# Early life tolerance depends on a subset of specialized dendritic cells and is reinforced by the skin microbiota

**DOI:** 10.1101/2022.06.23.497363

**Authors:** Antonin Weckel, Miqdad O. Dhariwala, Kevin Ly, Oluwasunmisola T. Ojewumi, Julianne B. Riggs, Jeanmarie R. Gonzalez, Laura R. Dwyer, Joy N. Okoro, John M. Leech, Margot S. Bacino, Grace D. Cho, Geil Merana, Niroshana Anandasabapathy, Yosuke Kumamoto, Tiffany C. Scharschmidt

## Abstract

Early life establishment of tolerance to commensal bacteria at barrier surfaces carries enduring implications for immune health but remains poorly understood. Here we show that this process is controlled by microbial interaction with a specialized subset of antigen presenting cells. More particularly, we identify CD301b^+^ type 2 conventional dendritic cells (DC) as a subset in neonatal skin specifically capable of uptake, presentation and generation of regulatory T cells (Tregs) to commensal antigens. In early life, CD301b^+^ DC2 are enriched for programs of phagocytosis and maturation, while also expressing tolerogenic markers. In both human and murine skin, these signatures were reinforced by microbial uptake. In contrast to their adult counterparts or other early life DC subsets, neonatal CD301b^+^ DC2 highly expressed the retinoic acid-producing enzyme, RALDH2, deletion of which limited commensal-specific Tregs. Thus, synergistic interactions between bacteria and a specialized DC subset critically support early life tolerance at the cutaneous interface.

## Introduction

Immune tolerance to environmental and commensal-derived antigens is fundamental to support both tissue and systemic immune homeostasis (Gensollen et al., 2016). Early life is recognized as a particularly important window for establishment of this tolerance, during which tissues are endowed with enriched tolerogenic capacity (Skevaki and Thornton, 2020). However, there are inherent obstacles to studying neonatal immunity, such as low cell numbers, small organism size, and limited access to human tissues. These have resulted in persistent knowledge gaps regarding the cell types and pathways instructing unique early life immune functions. Overcoming these challenges is critically important if we hope to not only treat established immune-mediated diseases but learn how to intervene early enough to prevent their onset and mitigate their impact.

Immune tolerance is especially important in skin, one of the body’s most expansive barrier interfaces (Gallo, 2017) that is subject to continuous environmental exposures and also home to a rich community of bacterial symbionts (Byrd et al., 2018). A growing body of work demonstrates that cutaneous bacteria function as key collaborators in many aspects of skin health. These commensals promote physical and antimicrobial barrier integrity, accelerate wound-healing, and tune adaptive immune cell function (Harris-Tryon and Grice, 2022). Skin is also a tissue frequently afflicted by allergic, autoimmune and other chronic inflammatory diseases. Failure to establish or maintain immune tolerance to skin commensal bacteria can incite or accelerate such disorders (Carmona-Cruz et al., 2022). Despite this, we still have very limited understanding of the cell types and mechanisms supporting immune tolerance to skin commensal bacteria.

We have shown that early life represents a preferential window for establishment of immune tolerance to skin commensal bacteria. As has been confirmed more recently in the gut (Akagbosu et al., 2022; Knoop et al., 2017; Zegarra-Ruiz et al., 2021), this tolerance is marked by generation of commensal-specific regulatory T cells (Tregs) that limit skin neutrophils upon later life bacterial re-exposure (Scharschmidt et al., 2015). It remains unclear, however, whether the preferential establishment of immune tolerance in early life reflects a global difference in regulatory capacity across the entire tissue or is directed by a specialized cell subset. The phenotype and function of antigen-presenting cells (APC), more specifically dendritic cells (DC) in neonatal skin represents an additional area of limited knowledge. Human fetal skin DCs have broadly been shown to possess tolerogenic function (McGovern et al., 2017), but whether such capacity is enriched in a defined cell subset and the mechanism linked to any such sub-specialization remain unknown. The major mechanisms by which neonatal DCs drive Treg generation in the cutaneous periphery and whether these are further conditioned by skin commensal bacteria are other outstanding questions.

Here, we took advantage of engineered bacterial strains, in vivo and ex vivo models of murine and human skin colonization, single-cell RNA sequencing and in vitro mechanistic studies to identify a functionally distinct subset of CD301b^+^ type 2 conventional DC (DC2) in neonatal skin. Enriched for phagocytic, maturation as well as tolerance markers, these cells were revealed to be uniquely equipped to uptake skin bacteria, present their antigens and generate commensal-specific Tregs. Notably, direct bacterial interaction reinforced these signatures. The regulatory capacity of neonatal CD301b^+^ DC2 was specifically tied to heightened retinoic acid production, implicating this as a neonatally-enriched tolerance mechanism. Collectively, these data reveal that early life tolerance at barrier surfaces does not reflect a broadly-expressed tissue feature, but rather a unique capacity of specialized cell subsets that is reinforced by the microbiota.

## Results

### CD301b^+^ DC2 support early life generation of commensal-specific Tregs

We have previously shown that neonatal skin colonization with *Staphylococcus epidermidis* engineered to express the model antigen 2w (*S. epi*-2w) supports a regulatory T cell (Treg) rich, antigen-specific CD4^+^ compartment (Leech et al., 2019). Moreover, when neonatally-colonized mice are re-challenged with *S. epi*-2w in the setting of skin abrasion they are more protected against skin inflammation than mice previously colonized during adult life (Scharschmidt et al., 2015). To uncover the mechanisms underlying the establishment of early life Treg to the microbiota colonization, we first compared the initial establishment of T cell responses to *S. epi*-2w association in neonatal versus adult mice. To this end, we performed repeated colonization of 7 day old (D7) pups or 6 week old adult mice and measured the antigen-specific and polyclonal CD4^+^ responses in the skin-draining lymph nodes (SDLN) 13 days later (Fig. 1A). Although the overall size of the 2w^+^CD44^+^CD4^+^ compartment was much smaller in neonatal than adult SDLN (Fig. S1A), the percentage of *S. epi*-specific Tregs was substantially increased in response to neonatal colonization (Fig. 1B). This was despite a slightly lower overall percentage of polyclonal Tregs in the SDLN at this age (Fig. S1B). This reinforces that early life establishment of long-term tolerance to skin commensal bacteria is closely linked to a Treg-rich primary antigen-specific response following initial *S. epi*-2w colonization.

**Figure 1:**
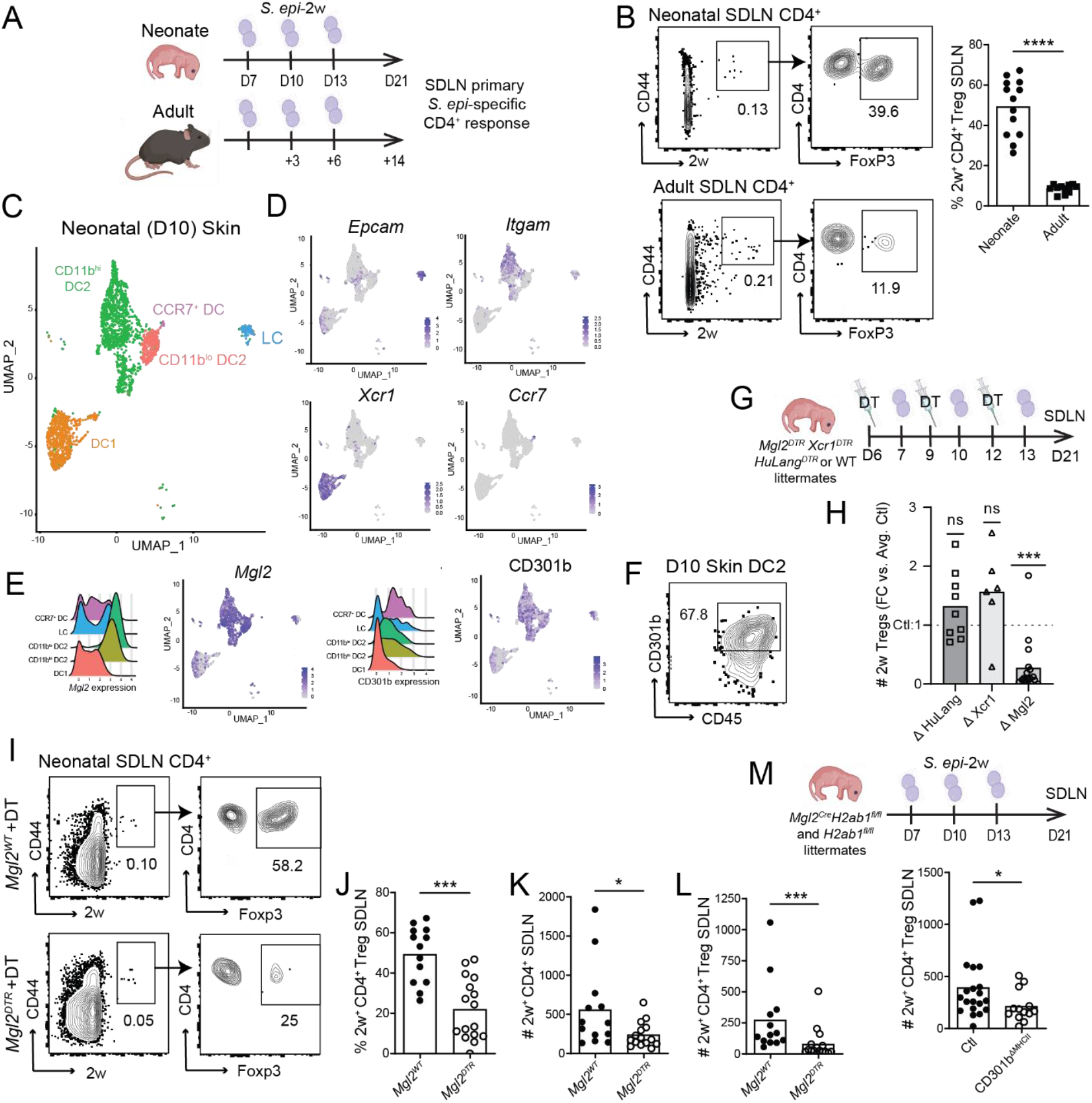
CD301b^+^ DC2 are required for early life generation of skin commensal-specific Tregs. (A) To measure the primary antigen-specific response to *S. epi*-2w in adult and neonatal mice, mice were colonized every 3 days for a total of 3 times and harvested a week following the last colonization; colonization was started on day 7 (D7) of life for neonates and on or after week 6 of life for adults. (B) Gating of tetramer 2w^+^CD44^+^ cells (pre-gated on live Lin(CD11b, CD11c, F4/80, B220)^neg^TCRβ^+^CD4^+^) from adult and neonatal skin draining lymph nodes (SDLN). Data from 2-3 independent experiments. (C) UMAP plot of major DC populations in D10 neonatal skin as revealed by scRNAseq on live MHCII^+^CD11c^+^ cells. (D) Feature plots of major genes used to identify skin DC subsets in (C). (E) Gene expression (left/center left) of *Mgl2* and its encoded protein CD301b (center right/right) in neonatal skin by CITE-seq. (F) Representative gates for CD301b expression in skin DC2 (pre-gated on live CD45^+^CD11c^+^MHCII^+^CD103^-^EpCam^-^) in a WT D11 mouse. (G) Experimental setup for (H-L). At D6, D9 and D12, neonatal mice were injected with DT and colonized 24h later with *S. epi*-2w, SDLN were harvested on D21. (H) Fold-change to littermate controls of total 2w^+^CD4^+^CD44^+^FoxP3^+^ cells in the indicated mouse models. One sample Wilcoxon test to hypothetical value of 1. (I) Gating of 2w^+^CD44^+^ cells (pre-gated on live Lin^neg^TCRβ^+^CD4^+^) of *Mgl2*^*WT*^ (top) and *Mgl2*^*DTR*^ (bottom) in D21 SDLN. (J) Percentage of Tregs in SDLN 2w^+^CD4^+^CD44^+.^ (K) Total 2w^+^CD4^+^CD44^+^ and (L) total 2w^+^CD4^+^CD44^+^FoxP3^+^ in SDLN. (M) Experimental set-up (top) and quantification (bottom). Total 2w^+^CD4^+^CD44^+^FoxP3^+^ in SDLN in of *Mgl2*^*cre*^*H2ab1*^*fl/fl*^ and *H2ab1*^*fl/fl*^ littermates. Pooled data from 3-4 independent experiments. Unless specified Mann-Whitney test was used with *p < 0.05, **p < 0.01, ***p < 0.001, ****p<0.0001, ns = not significant.

As dendritic cells shape the quality of CD4^+^ T cells responses to tissue antigens, we next sought to define the major DC subsets present in neonatal skin and SDLN by performing cellular indexing of transcriptomes and epitopes by sequencing (CITE-seq) on D10 mice. This confirmed *Epcam*^*hi*^ Langerhans cells (LC), *Xcr1*^*hi*^ CD103^hi^ type 1 conventional dendritic cells (DC1) and *Itgam*^*hi*^ and *Itgam*^*lo*^ type 2 conventional dendritic cells (CD11b^hi^ and CD11b^lo^ DC2s) as the four main skin DC types, with a separate small cluster of DCs that contain a mix of these markers but are defined primarily by their CCR7^+^ status (Figs. 1C-D, S1C). In the SDLN, CD11c^lo/hi^MHCII^hi^ migratory DCs (migDCs) were comprised of the same four DC subsets with different relative prevalence (Figs. S1D-E). Of note, the C-type lectin CD301b, encoded by *Mgl2*, is known to mark a portion of DC2 in adult murine skin and other tissues (Kumamoto et al., 2013; Linehan et al., 2015; Shin et al., 2016; Tatsumi et al., 2021). Our CITE-seq data confirmed enrichment of *Mgl2* and CD301b expression in both skin CD11b^hi^ DC2 and CD11b^lo^ DC2 clusters (Fig. 1E), with similar enrichment of CD301b protein but not RNA-level expression in SDLN migDC2 (Fig. S1F). Flow cytometry revealed that approximately two thirds of neonatal skin DC2s were CD301b^+^ by surface staining (Fig. 1F, S1G-I).

To systematically assess if a specific DC subset preferentially supports generation of Tregs specific for skin commensal antigens, we next used three complementary models to temporarily deplete each of the major skin DC subsets in the neonatal window during *S. epi*-2w colonization (Fig. 1G). As reported in adult mice (Yamazaki et al., 2013a), diptheria toxin (DT) administration to *Xcr1*^*DTR*^ mice led to preferential depletion of DC1 in the skin and SDLN (Fig. S1J). Likewise, Hu-Lang^DTR^ mice (Bobr et al., 2010) were a reliable model for LC depletion (Fig. S1K). No model yet exists for transient depletion of all DC2 (Loschko et al., 2016), thus we took an alternative strategy of using *Mgl2*^*DTR*^ mice to achieve 98% depletion of CD301b^+^ DC2 in the skin and SDLN (Fig. S1I and S1L). In this model, LC numbers were also somewhat reduced in neonatal skin but not neonatal SDLN (Fig. S1L), consistent with adult models and their expression of *Mgl2*/CD301b by CITE-seq (Fig. 1E, S1F). Examination of 2w^+^ CD44^+^ CD4^+^ T cells in the SDLN at D21 across these various models revealed a relative and absolute reduction in *S. epi-*specific Tregs only in *Mgl2*^*DTR*^ mice (Fig. 1H-J, S1M). Compared to DT-treated *Mgl2*^*WT*^ littermates, *Mgl2*^*DTR*^ mice demonstrated a reduction in the total number of *S. epi*-specific CD4^+^ T cells (Fig. 1K), which was driven entirely by a reduction of *S. epi-specific* Tregs (Fig. 1L) as *S. epi-*specific Foxp3^neg^ effector CD4^+^ T cell (Teffs) were unchanged (Fig. S1N). Polyclonal Treg percentages were equivalent between the groups (Fig. S1O), further emphasizing a specific defect in the *S. epi*-specific Treg compartment. Examination of *Mgl2*^*cre*^*H2ab1*^*fl/fl*^ (CD301b^ΔMHCII^) mice (Tatsumi et al., 2021), in which MHCII expression is reduced in CD301b-expressing cells (Fig. S1P), revealed a similar decrease in the number of *S. epi*-specific Tregs (Fig. 1M). This suggests that antigen-presentation by CD301b^+^ DC2 is specifically needed. Collectively, these results support a critical role for CD301b^+^ DC2 in neonatal generation of Tregs specific for skin commensal bacteria.

### Transient neonatal depletion of CD301b^+^ DC2 limits establishment of long-term tolerance to skin commensal bacteria

To assess the significance of neonatal CD301b^+^ DC2 in shaping the memory response to *S. epi*-2w, we used the *Mgl2*^*DTR*^ mouse system in our previously established re-challenge model (Fig. 2A). In this model, we previously showed that the absence of neonatal colonization with *S. epi*-2w leads to lack of immune tolerance in adulthood as evidenced by increased skin neutrophils upon *S. epi-*2w re-exposure in the context of epidermal abrasion (Scharschmidt et al., 2015). Here we colonized *Mgl2*^*WT*^ and *Mgl2*^*DTR*^ littermates with *S. epi*-2w during the first two weeks of life in tandem with DT administration to transiently deplete CD301b^+^ DC2. Two weeks later, upon recovery of CD301b^+^ DC2 in *Mgl2*^*DTR*^ (CD301b^ΔNeo^) mice (Fig. S2A-B), both groups were re-colonized with *S. epi*-2w in tandem with light tape-stripping (Fig. 2A). Post-challenge, CD301b^ΔNeo^ mice showed persistently reduced numbers and percentages of *S. epi-*specific Tregs in the SDLN (Fig. S2C).

**Figure 2:**
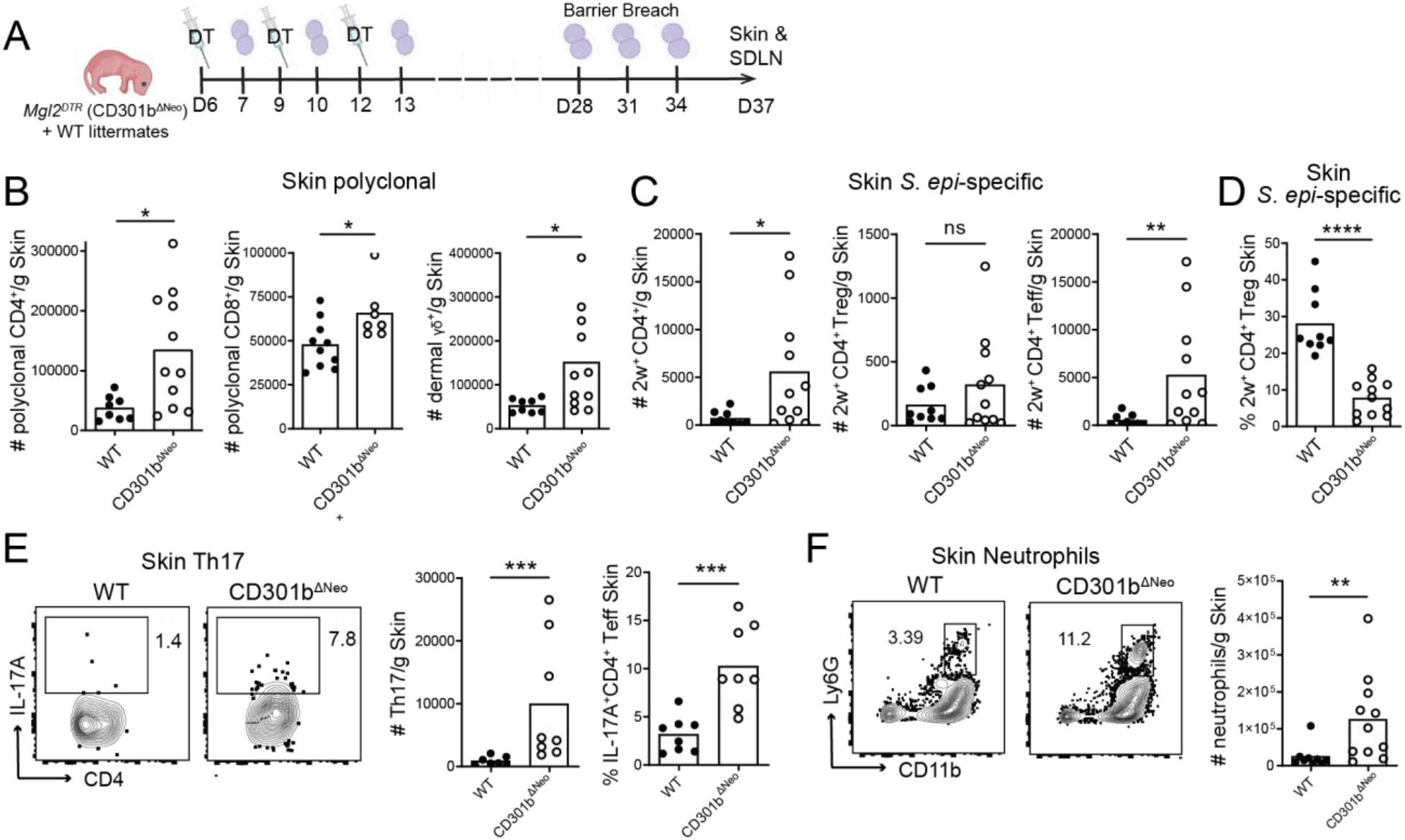
Neonatal CD301b^+^ DC2 are required for establishment of long-term tolerance to skin commensals. (A) To measure the long-term consequences of early life absence of CD301b^+^ cells, *Mgl2*^*wt*^ and *Mgl2*^*DTR*^ littermates treated neonatally with DT and *S. epi-*2w (as in Fig. 1G) were aged out for 2 weeks, then re-colonized every 3 days for a total of 3 times with accompanying epidermal barrier disruption by tape stripping. Mice were harvested 3 days after the last colonization (D37). (B) Total number of cells per gram of skin of polyclonal CD4^+^ (left), FoxP3^+^ CD4^+^ (center) and FoxP3^neg^ CD4^+^ (right). (C) Total number of cells per gram of skin of *S. epi*-specific 2w^+^CD4^+^CD44^+^ (left), 2w^+^CD4^+^CD44^+^FoxP3^+^ (center) and 2w^+^CD4^+^CD44^+^FoxP3^neg^ (right) in the skin. (D) Percentage of *S. epi*-specific Tregs in 2w^+^CD4^+^CD44^+^ in the skin. (E) Representative gating (pre-gated on live TCRβ^+^CD4^+^FoxP3^neg^) (left) and quantification of total number of cells per gram of skin of IL-17A^+^ CD4^+^ T effectors (center) and percentage of IL-17A^+^ CD4^+^ T effectors within the T effector population (right). (F) Representative gating (pre-gated on live CD45^+^CD3^neg^) (left) and quantification of neutrophils in skin (right). 2-3 independent experiments pooled. Unless specified, Mann-Whitney test was used with *p < 0.05, **p < 0.01, ***p < 0.001, ****p<0.0001, ns = not significant.

Transient neonatal removal of CD301b^+^ DC2 altered immune homeostasis of the skin with a significant increase in total numbers of CD4^+^ T cells, CD8^+^ T cells, and dermal *γδ* delta cells post-challenge (Fig. 2B). This accumulation of tissue lymphocytes was also reflected in higher numbers of *S. epi-specific* CD4^+^ T cells in skin (Fig. 2C), which was driven in large part by increased *S. epi-*specific Teffs (Fig. 2C). As in the SDLN, the percentage of *S. epi-*specific Tregs in skin was significantly reduced in CD301b^ΔNeo^ mice (Fig. 2D), despite equivalent polyclonal Treg percentages (Fig. S2D). These alterations in the commensal-specific response were accompanied by increased skin inflammation in CD301b^ΔNeo^ mice as evidenced by increased total numbers of IL-17 producing CD4^+^ (Th17) T cells (Fig. 2E) and neutrophils (Fig. 2F). These observations closely parallel the response to challenge seen in mice that were not colonized neonatally by *S. epi*-2w but only challenged as adults (Scharschmidt et al., 2015), suggesting that CD301b^+^ DC2 critically support establishment of long-term tolerance during neonatal colonization with commensal skin bacteria.

### Neonatal CD301b^+^ DC2 preferentially uptake skin bacterial antigens for presentation to CD4^+^ in the SDLN

Given the specific functional role for CD301b^+^ DC2 in shaping the neonatal CD4^+^ response to *S. epi-*2w, we next examined the capacity of the various skin DC subsets to uptake, carry to the SDLN, and present *S. epi-*expressed antigen. For this, we took first advantage of *S. epidermidis* engineered to express the phagolysosome resistant fluorophore, zsgreen (Matz et al., 1999) (*S. epi*-zsgreen) to track the dynamics of its in vivo uptake by DCs in neonatally colonized mice (Fig. 3A). This revealed preferential bacterial uptake by CD11b^hi^ DC2 in terms of both of the percentage of that subset loaded with *S. epi-*zsgreen (i.e. *S. epi-*zsgreen affinity) and the total number of loaded cells in the skin, both of which peaked 16 hours post-colonization (Fig. 3B-C). In the SDLN, the total number of migDCs increased between 8- and 16-hours post-colonization (Fig. S3A). Among these, CD11b^hi^ DC2 were again overrepresented in terms of their *S. epi-* zsgreen affinity and the total number of zsgreen^+^ cells (Fig. 3D-E), suggesting their transport of commensal antigen from skin to the SDLN. To understand if preferential uptake by DC2 was conserved across species we utilized a human neonatal foreskin model in which a physiological amount of *S. epi*-zsgreen was used to colonize the epidermal surface prior to examination of human skin DC subsets by flow cytometry 4 hours later (Fig. S3B-C). This revealed that human DC2 accounted for the majority of commensal-laden skin DCs (Fig. 3F-G).

**Figure 3:**
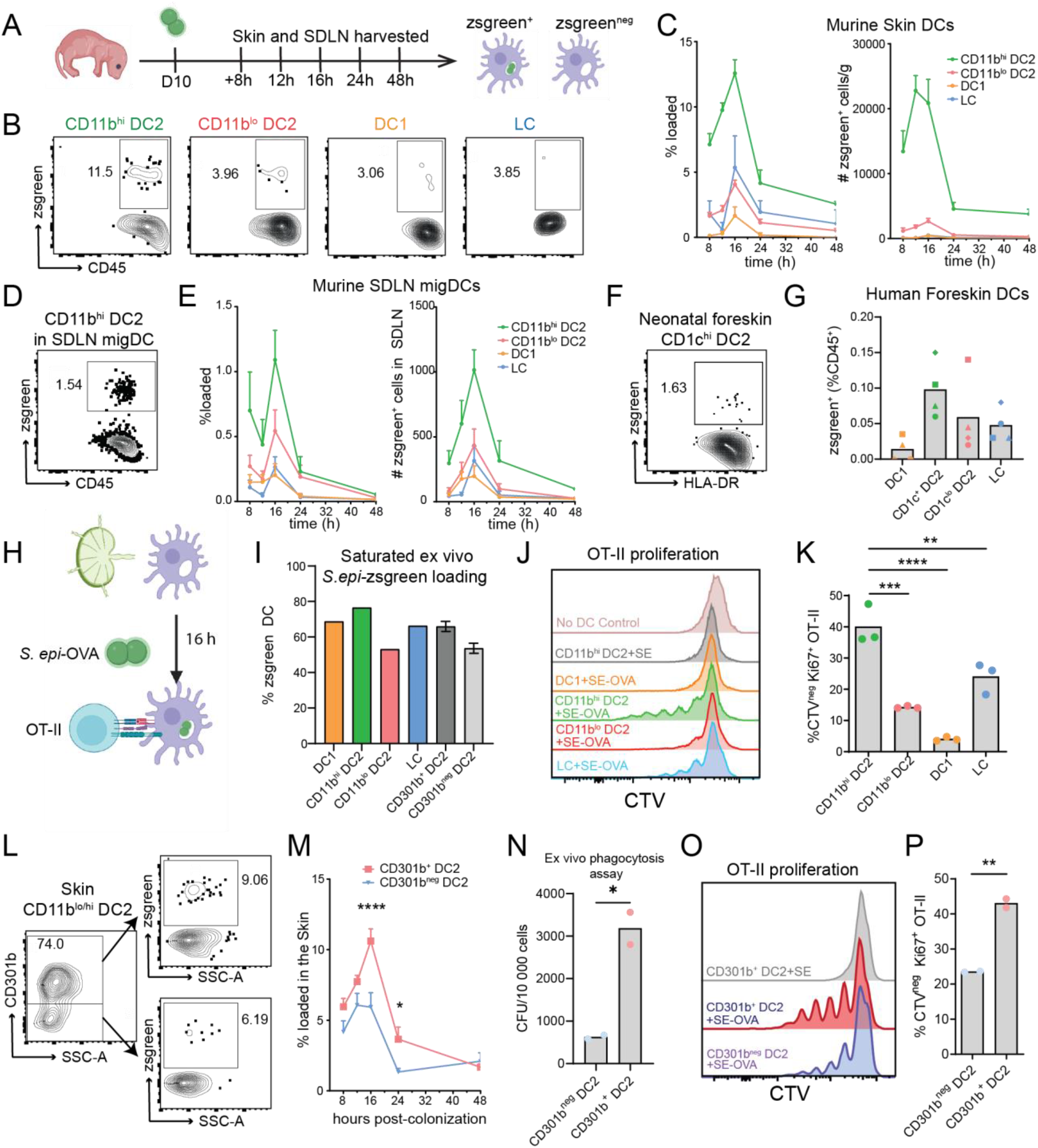
CD301b^+^ DC2s preferentially take up commensal bacteria in mouse and human neonatal skin to present these antigens in the SDLN. (A) Schematic related to B-E and L-M. D10 pups were colonized on skin with *S*.*epi*-zsgreen. Skin and SDLN were then harvested at multiple time points and DCs analyzed by flow cytometry; zsgreen^+^ DC correspond to those loaded with *S. epidermidis*. (B) Sample gating of zsgreen^+^ cells among skin DC subsets 16h after colonization (pre-gated as per Fig. S1G). (C) Time course of bacterial uptake by skin DC populations. Percentage of loaded cells (zsgreen^+^) as fraction of each population (left) and total number of loaded cells per gram of skin (right). (D) Sample gating of zsgreen^+^ cells in CD11b^hi^ migDC2 in the SDLN 16h post-colonization (pre-gated as per Fig. S1H). (E) Time course of zsgreen^+^ loaded migDC subsets in SDLN as a fraction of each population (left) or as total number (right). (F) Sample gating of bacterial uptake in human neonatal foreskin explants 4h after ex vivo colonization by *S. epi*-zsgreen (pre-gated as per Fig. S2C). (G) Uptake of commensal antigen by human neonatal foreskin DC populations 4h after colonization, as percentage of CD45^+^ cells. Each dot represents a separate human donor. (H) Schematic of the approach in I-K and N-P. Separate migDC subsets were sorted from the SDLN of D10 pups and incubated with *S. epi*-OVA for 2h, antibiotics were added to kill extracellular bacteria and then 16h later DCs were co-cultured with CTV-labelled OT-II CD4^+^ T cells for 72h prior to analysis of cell proliferation by flow cytometry. (I) Percentage of zsgreen^+^ cells by DC subset at the time T cells were added. One of three representative experiments is shown, SEM reflects technical replicates. (J) and (O) Histogram of OT-II CTV dilution, which is quantified in (K) and (P), illustrating different DC capacities to induce antigen-specific T cell proliferation after phagocytosing *S. epi*-OVA. Each dot represents a technical replicate from 1 of 3 representative experiments. (L) Sample gating and (M) time course of commensal antigen in neonatal skin by CD301b^+^ DC2 versus CD301b^neg^ DC2. (N) Ex vivo phagocytosis assay. Sorted DC from D10 SDLN were incubated for 2h with *S. epidermidis* at a MOI of 10, treated with gentamycin for 2h to kill adherent extracellular bacteria and plated to count intracellular CFUs, technical replicated of one experiment are shown. C, E and M SEM of 4 mice per time point. K One-way ANOVA. N, P students unpaired t-test. * p < 0.05, **p < 0.01, ***p < 0.001, ****p < 0.0001

Next, we examined the relative capacity of DC2 versus other subsets to present commensal antigen and induce antigen-specific CD4^+^ T cell proliferation. For this we engineered a strain of *S. epidermidis* to express the CD4^+^ epitope of ovalbumin (OVA), recognized by OT-II T cells (*S. epi-*OVA) (Merana et al., 2022).To test the capacity for antigen presentation independently from uptake, we sorted CD11b^hi^ DC2, CD11b^lo^ DC2, DC1 and LC from the migDC population of neonatal SDLN prior to directly incubating ex vivo overnight with *S. epi-*OVA (Fig. 3H). In this ex vivo approach, loading of the different DC subsets was similar (Fig. 3I). These *S. epi-*OVA-loaded migDCs were then co-cultured with cell trace violet (CTV)-labelled OT-II CD4^+^ T cells for 72 hours prior to assessing OT-II proliferation by flow cytometry. DCs loaded with non-OVA-expressing *S. epidermidis* and also wells with no DCs were used as controls. Although all DC subsets were capable of inducing proliferation of CD8^+^ T cells (Fig. S3D-E), these DC-T cell co-cultures revealed that CD11b^hi^ DC2, followed by LC and CD11b^lo^ DC2, were most capable of inducing proliferation in response to OVA and *S. epi*-OVA (Fig. 3J-K, S3F-G).

To determine if CD301b^+^ DC2 were specifically equipped to uptake skin bacterial antigens we re-examined our longitudinal data for *S. epi*-zsgreen loading of neonatal skin and SDLN DCs (Fig. 3A). In both skin DC2 and SDLN migDC2, this revealed a higher affinity for *S. epi*-zsgreen uptake among CD301b^+^ versus CD301b^neg^ cells (Fig. 3L-M, S3H-I). We confirmed this increased propensity for bacterial uptake using a separate ex vivo phagocytosis assay in which sorted CD301b^+^ or CD301b^neg^ DC2 from the SDLN were incubated with *S. epi*-zsgreen at a multiplicity of infection (MOI) of 10. This revealed three-fold more *S. epi* uptake by CD301b^+^ DC2 versus CD301b^neg^ DC2. Thus, in vivo differences in bacterial uptake can at least in part be explained by cell-intrinsic CD301b^+^ DC2 features (Fig. 3N). Using overnight ex vivo incubation of CD301b^+^ or CD301b^neg^ DC2 with *S. epi-*zsgreen at a MOI of 100 to normalize uptake (Fig. 3I), we then examined their relative capacity to present antigen to CD4^+^ T cells. Increased OT-II proliferation was induced by *S. epi*-OVA-loaded CD301b^+^ versus CD301b^neg^ DC2 (Fig. 3O-P), a difference which held also with OVA protein (Fig. S3J). Collectively these results indicate that DC2, specifically those that are CD301b^+^, are especially equipped for uptake of commensal bacterial antigens in the skin and their presentation to CD4^+^ T cells in the SDLN.

### CD301b marks a subset of activated DC2 enriched for phagocytic, maturation as well as regulatory markers

We next sought to more comprehensively identify defining features CD301b^+^ versus CD301b^neg^ DC2 in the skin and SDLN of neonates that might explain their unique role in the generation of commensal-specific Tregs. Focusing first on skin DC2, we returned to our CITE-seq data from D10 pups, now setting a threshold for *Mgl2*/CD301b expression to compare CD301b^+^ versus CD301b^neg^ DC2 cells (Fig. S1F). This revealed CD301b^+^ DC2 to be enriched for genes and gene pathways involved in antigen presentation and T cell activation together with genes coding for phagocytosis receptors (Fig. 4A-C, S4A, Table 1 & 2), which may explain their higher capacity to phagocytose and present bacterial antigens (Fig. 3). Skin CD301b^+^ DC2 expressed more CD80, CD86, CD40, OX-40L but also PD-L1, and PD-L2 as measured via spectral flow cytometry by percent of positive cells or average MFI (Fig. 4D-E), suggesting a mature but potentially also regulatory phenotype. Examination of SDLN CD301b^+^ versus CD301b^neg^ DC2 revealed transcriptional downregulation of phagocytic and other surface receptors as compared to the skin. However, SDLN CD301b^+^ DC2 retained higher protein level expression of CD80, CD86, CD40, PD-L1, and PD-L2 compared to CD301b^neg^ DC2 (Fig. S4B-C). They also showed increased RNA-level expression of *Aldh1a2*, which encodes the retinoic acid (RA) producing enzyme RALDH2 (Larange and Cheroutre, 2016) known to support Treg generation (Mucida et al., 2009), potentially suggesting further consolidation of their regulatory capacity in the SDLN (Table 3).

**Figure 4:**
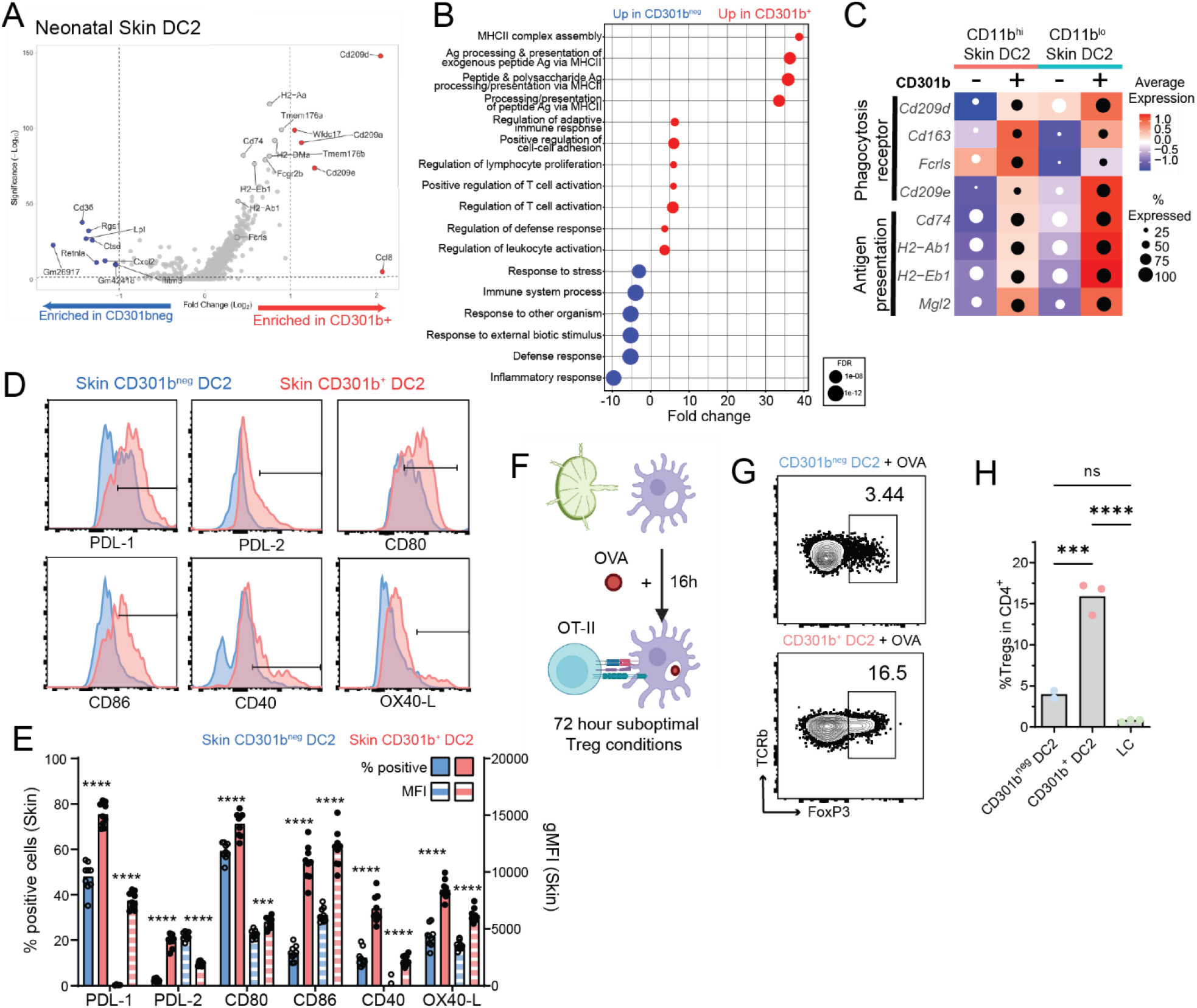
Expression of CD301b delineates a subset of DC2 enriched in phagocytic, maturation as well as regulatory markers. (A-C) derive from CITE-seq data introduced in Fig. 1C. (A) Volcano plot of top differentially expressed genes in CD301b^+^ versus CD301b^neg^ DC2 in D10 murine skin, complete list in Table 1 (B) Central functions enriched in each subset based on pathway analysis, complete list in Table 2 (C) Heat map of normalized average expression indicated genes in CD301b^+^ versus CD301b^neg^ cells in either CD11b^hi^ or CD11b^lo^ DC2 clusters and percentage of cells (circle size) in which expression was detected. All genes showed have a p-value <0.05 and were found in two independent scRNAseq experiments. (D-E) Spectral flow cytometry of activation markers in CD301b^+^ versus CD301b^neg^ DC2 from D10 skin 16h after colonization with *S. epidermidis*. (D) Representative histograms and gates used to delineate positive cells with corresponding (E) quantification of the % positive cells and geometrical mean of fluorescence intensity (gMFI) for the indicated markers. Each dot is biological replicate. 1 of 2 independent experiments are shown. Two-way ANOVA with Šídák’s multiple comparisons test. (F) Schematic of the DC-T cell assay used in (G) and (H) to measure DC capacity to promote Tregs. CD301b^+^ versus CD301b^neg^ DC2 were separately sorted from the migDC fraction of D10 SDLN and incubated with OVA protein overnight then with CTV-labelled OT-II T cells in the presence of IL-2 and TGF-β for 72h before flow cytometry to measure T cell proliferation. (G) Representative plots (pre-gated on Live CD4^+^TCRβ^+^). (H) Quantification of one of three representative experiments with technical replicates shown. One-way ANOVA. *p < 0.05, **p < 0.01, ***p < 0.001, ****p<0.0001.

Based on the in vivo data from *Mgl2*^*DTR*^ mice and the mature and regulatory phenotype of CD301b^+^ DC2, we explored their functional capacity to support Treg generation ex vivo. CD301b^+^ and CD301b^neg^ DC2 were sorted from the SDLN and incubated overnight with OVA protein before co-culture with OT-II CD4^+^ under suboptimal Treg-inducing conditions (Brown et al., 2019a) (Fig. 4F). This revealed a preferential capacity for Treg generation by CD301b+ versus CD301b^neg^ DC2 (Fig. 4G-H). Additionally, while LC were able to support CD4^+^ T cell proliferation at levels comparable to DC2 (Fig. S3F-G), they did not support Treg generation in this ex vivo assay. Collectively, this supports an intrinsic preferential capacity for antigen uptake by CD301b^+^ DC2 in skin as well as for antigen presentation and Treg generation in the SDLN.

### Uptake of skin commensal bacteria sustains maturation as well as tolerogenic features of CD301b^+^ DC2

The context in which a DC encounters an antigen, i.e. with or without certain pathogen-associated or damage-associated molecular patterns (PAMP/DAMPs), influences the secondary and tertiary signals presented by that DC in tandem with antigen. These DC-expressed co-stimulatory surface molecules and secreted cytokines, in turn, help dictate the phenotype and function of antigen-specific T cells primed by that DC. Our profiling of CD301b^+^ DC2 in neonatal skin and SDLN suggested that under homeostatic conditions that these cells have a mature but also tolerogenic phenotype, capable of supporting Tregs. To understand how direct encounter with skin commensal bacteria and uptake of *S. epidermidis* antigen further impact this phenotype, we again took advantage of our ability to identify and study skin DCs that phagocytose *S. epi-*zsgreen. After skin colonization of neonatal mice, we separately sorted zsgreen^+^ and zsgreen^neg^ skin DCs for analysis by single cell RNA sequencing (scRNAseq) (Fig. 5A). This revealed *S. epidermidis* uptake by skin CD301b^+^ DC2 to be correlated with higher expression of genes known to mediate regulatory DC functions, including Treg generation: *Cd80/Cd86* (Bar-On et al., 2011; Francisco et al., 2009; Salomon et al., 2000), *Fas* (Qian et al., 2013), *Cd274* (Francisco et al., 2009; Vogel et al., 2022) and *Socs1* (Fu et al., 2009) (Fig. 5B). Transcript levels of the Treg-promoting cytokine, *Il10* (Chaudhry et al., 2011), were also increased in zsgreen^+^ CD301b^+^ DC2, whereas those for *Il1b* and *IL-6*, which promote Th17 responses, were expressed more in zsgreen^neg^ CD301b^+^ DC2. For most of these markers, protein level increases in zsgreen^+^ versus zsgreen^neg^ skin CD301b^+^ DC2 from both the skin and SDLN were confirmed by spectral flow cytometry (Fig. 5C, S5A-B). Some additional markers, e.g. OX40L (Merana et al., 2022), were additionally found to be increased in *S. epi-*zsgreen-containing CD301b^+^ DC2 at the protein but not mRNA level (Fig. 5C. S5A-B).

**Figure 5:**
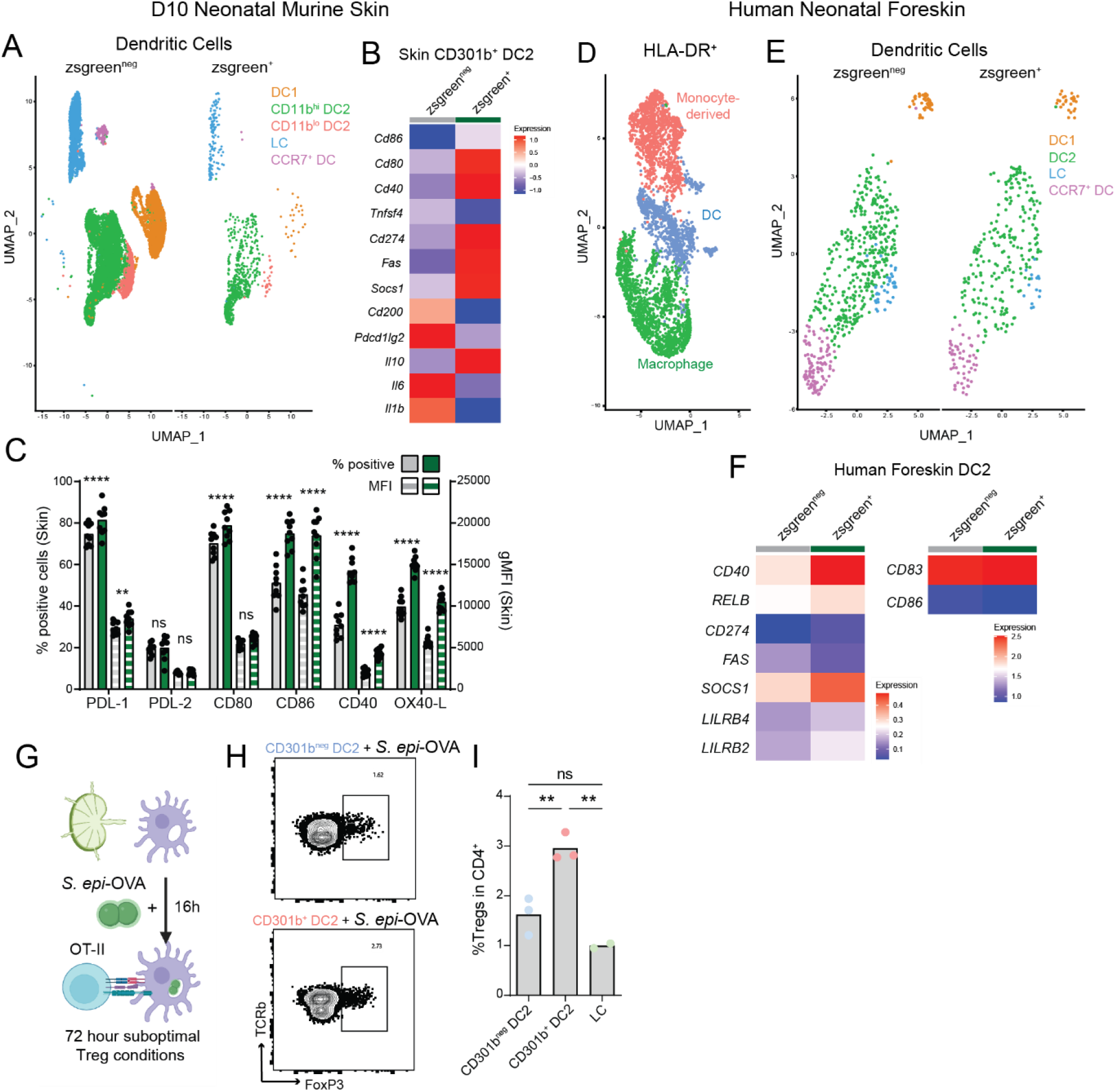
Uptake of commensal bacteria reinforces maturation and tolerogenic features of murine and human neonatal skin DC2. (A-B) D10 neonatal mice were colonized with *S*.*epi*-zsgreen and skin was harvested 16h later, with commensal antigen-loaded (zsgreen^+^) and unloaded (zsgreen^neg^) DCs separately sorts and submitted for scRNAseq. (A) UMAP of zsgreen^+^ vs zsgreen^neg^ DCs and (B) heatmaps comparing expression of select differentially genes between zsgreen^+^ vs zsgreen^neg^ CD301b^+^ DC2 (CD301b status defined based *Mgl2* expression). (C) Spectral flow cytometry comparing protein-level expression of key markers zsgreen^+^ vs zsgreen^neg^ CD301b^+^ DC2 in neonatal skin 16h after colonization with *S. epi-*zsgreen colonization. Quantification of the % positive cells and gMFI for the indicated markers in loaded vs non loaded CD301b^+^ DC2. Each dot is a biological replicate. 1 of 2 independent experiments are shown. Two-way ANOVA with Šídák’s multiple comparisons test. (D-F) Human neonatal foreskin was incubated 4 h with *S. epi*-zsgreen as detailed in Fig. S3B and loaded (zsgreen^+^) versus non-loaded (zsgreen^neg^) live CD45^+^ CD16^neg^ HLADR^+^ cells were separately sorted and submitted for scRNAseq. (D) UMAP of all cells combined, (E) UMAP of DC clusters split by zsgreen status, and (F) heat maps comparing expression of select genes between zsgreen^+^ and zsgreen^neg^ DC2 in human foreskin. (G) Schematic of the DC-T cell assay used in (H) and (I). Indicated DC subsets were sorted from the migDC fraction of D10 SDLN, incubated with *S. epi*-OVA overnight and then with CTV-labeled OT-II T cells for 72h in the presence of IL-2 and TGF-β. (H) Representative plots (pre-gated on Live CD4^+^TCRβ^+^) of showing percentage of OT-II Tregs and (I) and quantification thereof. Technical replicates are shown from one of 3 replicate experiments with similar results. One-Way ANOVA. *p < 0.05, **p < 0.01, ***p < 0.001, ****p<0.0001, ns = not significant.

Taking advantage of our neonatal foreskin explant system, in which we validated DC2 uptake of *S. epi*-zsgreen (Fig. 3), we assessed if *S. epidermidis* phagocytosis similarly polarized human skin DCs. After 4 hours of *S. epi-*zsgreen colonization, zsgreen^+^ and zsgreen^neg^ HLA-DR^+^ cells were separately sorted and submitted for scRNAseq, after which clusters of dendritic cells subsets were identified using classical markers (Figs. 5D-E, S5C-D) (Haniffa et al., 2015). Despite the short period post-colonization, *S. epi-*zsgreen-loaded DC2 demonstrated higher expression of many parallel markers of human regulatory DC function (Fig. 5F) (Domogalla et al., 2017).

To assess if CD301b^+^ DC2 capacity to support regulatory T cells was sustained upon *S. epidermidis* uptake, we returned to our ex vivo DC-T cell assay. This time, CD301b^+^ vs CD301^neg^ DC2 were incubated overnight with *S. epi-*OVA prior to co-culture with OT-II under sub-optimal Treg conditions (Fig. 5G). Only CD301b^+^ DC2 were capable of generating a significant amount of Tregs (Figs. 5H-I). In contrast, LC and CD301^neg^ DC2 which were able to generate significant proliferation of OT-II in response to *S. epi-*OVA (Fig. 3J-P), did not demonstrate comparable capacity to support *S. epi-*specific Tregs (Figs. 5H-I). Taken together, these data indicate that the a priori tolerogenic capacity of CD301b^+^ DC2 is maintained upon uptake of skin commensal bacteria, consistent with the in vivo evidence (Figs. 1, S1) that they promote commensal-specific Tregs.

### RALDH2 activity, enriched in neonatal CD301b^+^ DC2, supports early life commensal-specific Tregs

Although the mechanisms by which DCs promote Treg generation are multiple and complex (Steinman et al., 2003), DC production of retinoic acid (RA) has been linked to this function by multiple groups (Coombes et al., 2007; Mucida et al., 2009). This is especially true in gut-associated lymphoid tissue (Coombes et al., 2007; Sun et al., 2007), where the RA-precursor, Vitamin A, is in high abundance from dietary sources. Whether there is role for RA in tolerance at the cutaneous interface, if this tolerance mechanism differs by age and if it can be influenced by commensals remain unanswered questions. DC production of RA is controlled by retinaldehyde dehydrogenase 2 (RALDH2), encoded by *Aldh1a2* (Guilliams et al., 2010). Our scRNAseq identified increased A*ldh1a2* expression in CD11b^+^ and CD301b^+^ migDC2 versus other subsets (Fig. 6A, Table 3). A two-to three-fold increase was confirmed by qRT-PCR on CD301b^+^ and CD301b^neg^ migDC2 sorted from neonatal SDLN (Fig. 6B and Fig. S6A). Parallel analysis of adult SDLN DCs illustrated high A*ldh1a2* expression to be a specific feature of neonatal CD301b^+^ DC2 versus their adult counterparts (Fig. 6B and Fig. S6A). Increased protein-level expression of RALDH was likewise seen in neonatal CD301b^+^ DC2 (Fig. 6C and Fig. S6B). To understand if commensal uptake influences RALDH activity in CD301b^+^ DC2, we measured enzyme levels in sorted cells with or without ex vivo phagocytosis of *S. epi-*zsgreen. This revealed increased RALDH activity in zsgreen^+^ versus zsgreen^neg^ CD301b^+^ DC2 (Fig. 6D), suggesting that skin commensals reinforce this key tolerogenic pathway.

**Figure 6:**
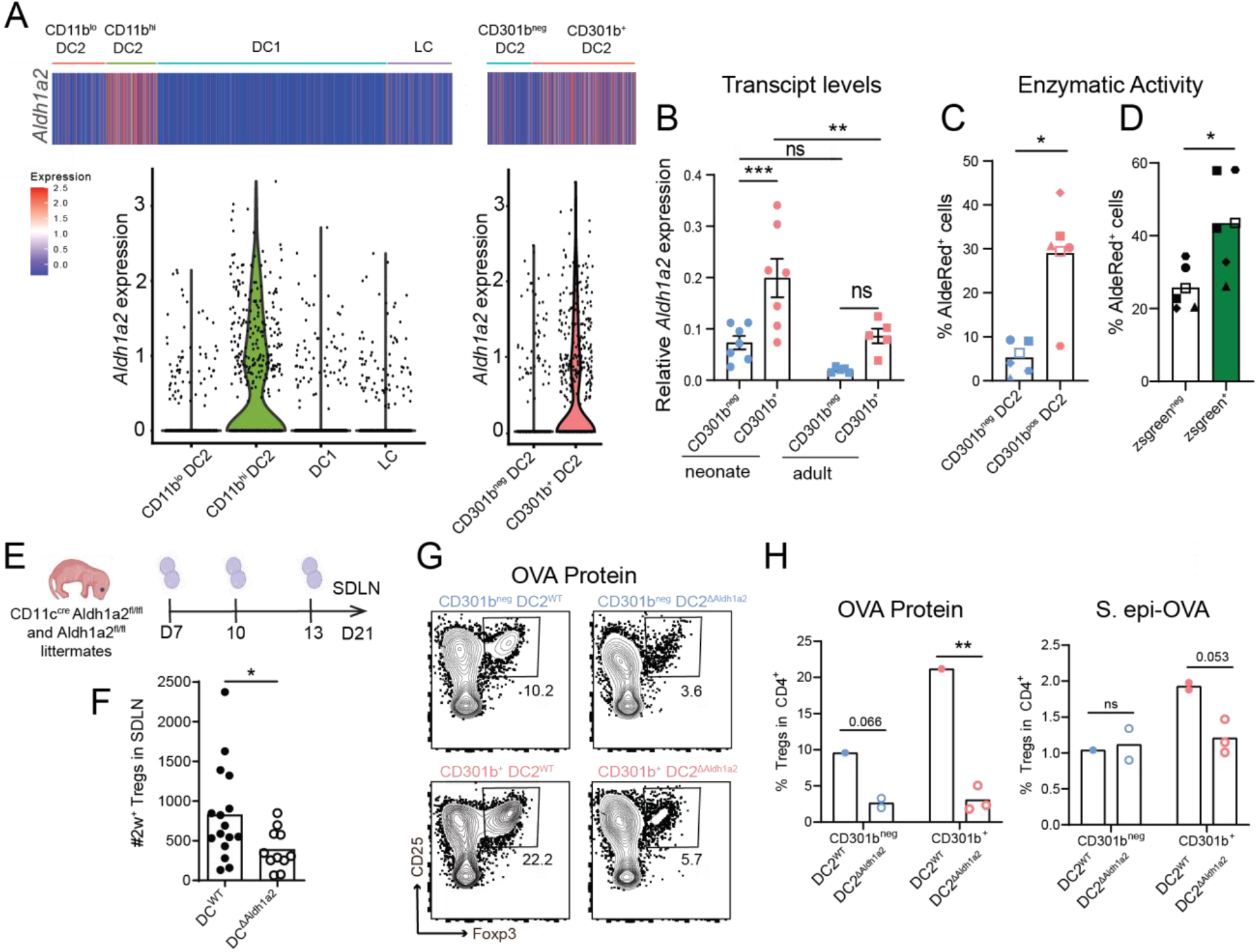
*Aldh1a2* expression is enriched in neonatal CD301b^+^ DC2s and augmented by *S. epidermidis* uptake thereby promoting generation of commensal-specific Tregs. (A) Heatmap (top) and Violin Plot (bottom) of *Aldh1a2* expression in subpopulations of D10 SDLN migDC. (B) qRT-PCR of *Aldh1a2* expression in CD301b^+^ versus CD301b^neg^ migDC2 sorted from SDLN of D10 mice and adult mice. Each dot is an independent mouse, Two-way ANOVA. (C) ALDH activity in CD301b^+^ versus CD301b^neg^ DC2 sorted from SDLN. (D) ALDH activity in zsgreen^+^ versus zsgreen^neg^ CD301b^+^ DC2 after a 16h incubation of DCs in the presence of *S. epidermidis-*zsgreen. Each dot represents a biological replicate of cells pooled from two mice. (E) Experimental setup for (F). (F) Total number of 2w^+^CD4^+^CD44^+^ FoxP3^+^ *S. epi-*2w-specific Tregs in D21 SDLN of neonatally-colonized *Cd11c*^cre^ *Aldh1a2*^fl/fl^ and *Aldh1a2*^fl/fl^ control littermates. Mann-Whitney test. (G-H) CD301b^+^ and CD301b^neg^ DC2 from the SDLN of neonatal *Cd11c*^cre^ *Aldh1a2*^fl/fl^ and *Aldh1a2*^fl/fl^ mice were incubated with OVA or *S. epi*-OVA followed by 72h co-culture with CTV-labelled OT-II in suboptimal Treg-inducing conditions. (G) Representative plots (pre-gated on Live CD4^+^TCRβ^+^) and (H) quantification of OT-II Tregs. One of two representative independent experiments is shown. Dots indicate technical replicates from DCs sorted from pooled neonates. Wilcoxon test. *p < 0.05, **p < 0.01, ***p < 0.001, ns = not significant.

To understand the in vivo contribution of DC-derived RADLH2 in antigen-specific Treg generation to skin commensal bacteria we returned to our antigen-specific model. *Cd11c*^*cre*^ *Aldh1a2*^*fl/fl*^ (DC^ΔAldh1a2^) neonates were colonized alongside control *Aldh1a2*^*fl/fl*^ (DC^ΔWT^) littermates with *S. epi-*2w on days 7, 10 and 13 of life, prior to harvest of SDLN at weaning age (Fig. 6E). This revealed a significant reduction in the number of *S. epi-*2w Tregs generated in the absence of RALDH2 expression by CD11c^+^ DCs (Fig. 6F). To probe the functional importance of RA production more specifically in CD301b^+^ DC2 versus other subsets, we isolated CD301b^+^ versus CD301b^neg^ DC2 from the SDLN of DC^ΔAldh1a2^ versus DC^WT^ mice and performed DC-T cell co-cultures under sub-optimal Treg conditions. This revealed a significant contribution of RALDH in Treg generation (Fig. 6G-I), but not T cell proliferation (Fig. S6C), by CD301b^+^ DC2 in response to both soluble and commensal-expressed OVA antigen. *Aldh1a2* deletion in CD301b^neg^ DC2 also led to a decrease in Treg generation to soluble protein but not in response to *S*.*epi*-OVA (Fig. 6G-I). Thus, RA production by neonatal CD301b^+^ DC2, which is enriched versus other subsets or their adult counterparts, plays a key role in their preferential capacity to support generation of commensal-specific Tregs.

## Discussion

Here we demonstrate CD301b^+^ DC2 as a critical cell population that supports early life establishment of peripheral tolerance to skin commensal bacteria. This function stems in part from a preferential capacity to sample bacterial antigens, a feature shared with DC2 in human neonatal skin. However, while other DC subsets, such as LC, can also efficiently uptake skin bacteria and present commensal antigens to induce CD4^+^ T cell proliferation, CD301b^+^ DC2 demonstrate a unique propensity for Treg generation. Uptake of commensal bacteria by human or murine DC2 further reinforces key features of their genetic program, namely maturation and antigen presentation. Moreover, it sustains CD301b^+^ DC2 expression of tolerogenic markers and augments their production of RALDH, which we show is a key mechanism by which these cells support Treg generation. Collectively, these results elucidate key cellular and molecular mechanisms supporting early life tolerance to commensals at the cutaneous interface.

In contrast to the skin, the intestines contain specialized cells for uptake of luminal antigens. These goblet cells (GC) have been shown to facilitate uptake of antigens from commensal bacteria in the pre-weaning window, forming a cooperative relationship with the DC subsets that present these antigens (Kulkarni et al., 2019; Rios et al., 2015). Our work demonstrates evolution in skin of parallel but distinct mechanisms to support effective sampling of bacterial antigens during early life, namely a DC population especially equipped with this capacity. Additional work is needed to define the specific tissue niche of skin DC2s and how this might facilitate their capacity for bacterial uptake. However, our ex vivo phagocytosis assays indicate this behavior derives, at least in part, from cell-intrinsic features, such as expression of microbial-associated molecular pattern (MAMP) receptors and phagocytic receptors. In the intestine, CD103^+^ DCs are recognized as a key DC subset supporting tolerance to commensal bacteria via peripheral induction of Tregs (Coombes et al., 2007). More recently, Aire-expressing DCs in the intestinal lymph nodes have been identified as an important cell type contributing to this function (Akagbosu et al., 2022; Wang et al., 2021). While further investigation is needed to understand potential overlapping features and functions between CD301b^+^ DC2 and these DC subsets, initial evidence supports that there is some degree of barrier site-specific restriction of this function to distinct DC types.

CD301b^+^ DC2 have been implicated in supporting a range of different types of T cell responses. They have been studied most in depth in the context of allergic inflammation where they facilitate differentiation of type 2 helper (Th2) cells (Kumamoto et al., 2013). They can also optimize priming of naïve CD4^+^ T cells in lymph nodes by detaining these cells near the high endothelial venule for maximal antigen scanning (Tatsumi et al., 2021). In other models, they have been shown to support Th17 responses (Kim et al., 2018; Linehan et al., 2015) and to promote clonal deletion of thymic T cells (Breed et al., 2022). This suggests that the function of CD301b^+^ DC2 is contextual, with our findings reflecting their contributions in a setting of early life tissue homeostasis.

Our data, consistent with that from thymic CD301b^+^ DC2 (Breed et al., 2022), indicates maturation and antigen presentation capacity as features strongly associated with DC2 expression of CD301b. While in certain contexts this state may primarily support Th2/Th17 differentiation, we observed concurrent upregulation of molecules associated with regulatory DC function, as has been observed in tumor-associated ‘mReg’ DC (Maier et al., 2020) and migratory DCs more generally (Nirschl et al., 2017). Notably, we did not find DCs capacity to induce commensal-specific T cell proliferation to be strictly coupled with their ability to promote Tregs. LCs efficiently stimulated CD4^+^ T cell proliferation but did not Tregs to *S. epi*-OVA, whereas the absence of RALDH in CD301b^+^ DC2 reduced Treg generation without affecting CD4^+^ proliferation. Collectively, these studies strengthen a narrative that the capability of DCs to induce Tregs is not necessarily linked to a state of relative immaturity.

Other recent work examining DC2 in the thymus (Breed et al., 2022) has highlighted CD301b as marking a subset of DC2 with distinct functional capacity. Our results suggest that that specific functional capacity differs by tissue. We show in skin that expression of CD301b in DC2 strongly correlates with that of other phagocytic receptors and that these cells are efficient at bacterial uptake. Type 2 cytokines have been shown to promote CD301b^+^ DC2 in the thymus (Breed et al., 2022) and CD11b^lo^ DC in the skin (Mayer et al., 2021), but we found CD301b^+^ DC2 cells still present in the skin of germ-free as well as *Il13*^*-/-*^*Il4r*^*-/-*^ neonates (data not shown). Thus, the specific pathways driving expression of CD301b and associated programs in skin DC2 remain to be determined.

The ability of DCs to support Treg induction and proliferation has been tied to many different pathways and molecules (Domogalla et al., 2017), many of which are expressed by CD301b^+^ DC2. Synergy across multiple tolerogenic mechanisms likely supports Treg generation by these and other tolerogenic DC populations. We were intrigued to find, however, that the RALDH pathway for RA production was specifically enriched among CD301b^+^ DC2 and increased upon uptake of *S. epidermidis*. RA production is known to support peripheral Treg induction and limit Th1/Th17 differentiation via effects on TGFβ signaling (Mucida et al., 2007; Xiao et al., 2008). Until now, it has been described primarily in gut-associated lymphoid tissues (Sun et al., 2007), specifically CD103^+^ DC (Coombes et al., 2007), and has also been implicated specifically in generation of alpha4 beta 7 integrin+ gut-homing T cells (Benson et al., 2007). While RA production capacity had been hinted at previously in subsets of SDLN DC (Guilliams et al., 2010), our work clearly pinpoints this as a specific feature of CD301b^+^ DC2 and one that is preferentially enhanced in early life. Uptake of *S. epidermidis* was able to further increase RALDH activity/expression in these cells, consistent with prior work suggesting that TLR2 and Myd88 signaling can provide positive feedback loops for RA production (Villablanca et al., 2011; Wang et al., 2011). Further work is needed however to tease apart interactions between skin commensal bacteria and cutaneous DCs and how these influence CD301b^+^ DC function.

There is growing recognition that early life microbial interactions impact long-term disease predisposition (Brodin, 2022). Intervening therapeutically or preventatively in this window will require deeper understanding of key host-microbe mechanisms active in this early period. This will include identifying microbial species and molecules that carry harmful or helpful effects. Equally important will be dissecting core host pathways that augment or impair tolerance. Cutaneous as well as orally-directed therapies represent core avenues for potential intervention (Brough et al., 2022). In this context, our work sheds light on the importance of dissecting tissue-specific mechanisms that facilitate early life tolerance to commensal and other environmental antigens.

## Acknowledgements

We thank James Moon for 2w tetramer reagents and expertise, Yasmine Belkaid Ph.D., James Gardner M.D., Ph.D., Margaret Lowe Ph.D., Clifford Lowell M.D., Michael Rosenblum M.D., Ph.D., and Ari Molofsky M.D., Ph.D. for helpful discussions. We thank Michelle Chu for help in cloning the *S. epi*-OVA strain, Yongmei Hu for mouse husbandry, and Sepideh Nozzari for genotyping. We appreciate sharing of the *huLang*^*DTR*^ mice by Daniel Kaplan M.D. Ph.D., the *Xcr1*^*DTR*^ mice by Max Krummel Ph.D., and the *Raldh2*^*fl*/fl^ mice by Randolph Noelle Ph.D. We acknowledge UCSF’s PFCC (RRID:SCR_018206) for assistance in flow cytometry and cell sorting, supported by NIH grants P30 DK063720 and 1S10OD021822-01. We acknowledge the UCSF Genomics CoLab and its staff members in supporting generation of CITE-seq and scRNAseq data. M.O.D. is supported by NIAMS K99AR079554. This work was primarily funded by T.C.S. grants NIAMS R01AR080034, NIAID DP2AI144968, and Burroughs Wellcome Fund CAMS-1015631.

## Author Contributions

A.W. and T.C.S. designed the studies and wrote the manuscript. A.W. performed the experiments and analyzed the data. M.O.D., K.L., O.T.O, J.B.R., J.R.G, L.R.D, J.N.O., J.M.L, M.S.B, G.D.C. and G.M. assisted with experiments. Y.K, and N.A. provided resources and input on experimental design. T.C.S. oversaw all study design and data analysis. All authors discussed results and commented on the manuscript.

## Experimental Model and Subject Details

### Experimental Animals

Wild-type C57BL/6 mice were originally purchased from Jackson Laboratories (Bar Harbor, ME), then bred and maintained in the UCSF specific pathogen-free (SPF) facility on the Parnassus campus for use in experiments., *Xcr1*^*DTR-Venus*^ mice were purchased from Jackson (Yamazaki et al., 2013b) and bred in-house. *Mgl2*^*DTR-eGFP*^ and *Mgl2*^*Cre*^*H2ab1*^*fl/fl*^ were a gift from Yosuke Kumamoto and Akiko Iwasaki (Kumamoto et al., 2013; Tatsumi et al., 2021). *Cd11c*^*Cre*^ *Aldh1a2*^*fl/fl*^ mice (Ji et al., 2006)were a gift from Elizabeth Nowak (Noelle Lab, Dartmouth University), HuLang^DTR^ (Bobr et al., 2010) mice were gift from Daniel Kaplan (University of Pittsburg).

All animals were 7 days to 10 weeks old at the time of experiments. Littermates of the same sex were socially housed under a 12 h light/dark cycle and randomly assigned to experimental groups whenever possible. Animal work was performed in accordance with the NIH Guide for the Care and Use of Laboratory Animals and the guidelines of the Laboratory Animal Resource Center and Institutional Animal Care and Use Committee of the University of California, San Francisco.

### Bacterial Strains and Culture Conditions

*Staphylococcus epidermidis* (*S. epi*) strain Tü3298 (Allgaier et al., 1986; Augustin and Gotz, 1990) was used in this study and grown in tryptic soy broth at 37ºC. Bacterial media was supplemented with 10 µg/mL erythromycin for plasmid selection. In current and published work, *S. epi* has been engineered to express the 2w model antigen linked to the fluorophore mCherry and zsgreen under control of the *agr* promoter via plasmid pJL74-2W-gpmCherry and pJL74-2W-zsgreen (Leech et al., 2019). In this work, the same Tü3298 strain was engineered to express the OVA peptide antigen via modification of the original pJL74-2W-gpmCherry plasmid.

## Method Details

### Bacterial Skin Colonization and Light Skin Abrasion Models

*S. epi*-2w was cultured for 48 hours to achieve high 2w-mCherry expression as measured by flow cytometry, then washed and re-suspended in PBS to obtain 10^8^-10^9^ colony-forming units (CFUs) at a volume of 100 μL per mouse. *S. epi*-2W was then applied via a plastic pipette and a sterile PBS-soaked cotton-tipped swab to the back skin of mice on days 7, 10, and 13 for neonatal colonization. To mimic physiologic exposure of mice to skin *S. epi*-2w in the context of light skin abrasion during adulthood, back hair was first removed using small animal clippers and depilatory cream (Nair™ Hair Remover Body Cream) on day 28, followed by repeated application and removal of adhesive tape on days 28, 31, and 34 (Shurtape HP-500). Tissues were harvested on day 37.

### Diptheria Toxin Injection

Mice were intraperitoneally injected with 25 ng.g^−1^ body weight DT (Sigma, D0564) dissolved in PBS at 2.5 ng.µl^−1^ concentration.

### APC-T cell In Vitro Assay

The APC-T cell assay using OVA protein and bacteria was informed by previous work (Brown et al., 2019b), with modifications specific to our model.

#### APCs

Skin-draining lymph nodes were harvested and processed over sterile 100 μm cell strainers in 1 mL of T cell media (RPMI supplemented with HyClone Characterized Fetal Bovine Serum (FBS), 1% penicillin-streptomycin, β-mercaptoethanol, HEPES and GlutaMAX™). Lymph nodes were transferred to digestion media (HBSS, 5% Fetal Calf Serum, 1% HEPES) containing 500U/mL of Collagenase D (Sigma-Aldrich, Cat 11088866001) and 20 mg/mL of DNAse I, mechanically disrupted and incubated at 37°C for 25 min. Organs were resuspended with a glass pipet and EDTA (10 mM final) was added, and organs were incubated another 5 minutes at 37°C. Tissues were then processed through a 100 μm cell strainers, washed and stained. Dendritic cells were sorted on a BD FACSAriaII using a 100-µm nozzle and sorted into RPMI medium supplemented with 10% FBS. 10 000 Dendritic cells were plated in 100 μL of T cell medium with no antibiotics.

#### Antigens

Soluble antigen: OVA protein was added at 0.5 mg/mL to sorted DC in T cell medium overnight. *Bacteria*: 48 hours cultures of *S*.*epi-*OVA were spun down, washed and resuspend in T cell medium with no antibiotics. Bacteria were added to sorted DCs to a multiplicity of infection of 100 (10^6^ bacteria per well), then incubated for 2 hours at 37ºC. Gentamycin was added at 25 µg/mL for 2 hours. Cells were washed and incubated overnight at 37ºC in T cell medium with antibiotics.

#### T cells

SDLN and spleen of adult OTII mice were harvested and processed over sterile wire mesh in 2 mL of complete RPMI media before cell isolation. Cells were ACK lysed to remove red blood cells and CD4^+^ T cells were isolated via EasySep™ Mouse CD4+ T Cell Isolation Kit (Catalog No. 19852). Isolation efficiency was verified via flow cytometry. CD4^+^ T cells were then labeled with the CellTrace™ Violet Cell Proliferation Kit (Invitrogen™, Catalog No. C34557) before co-culturing at 37ºC with antigen-pulsed dendritic cells at a ratio of 1:10. For suboptimal Treg inducing conditions, IL2 (200 U/mL, GoldBio, # 1310-02-100) and TGFβ (2 ng/mL, Peprotech, #100-21C-10UG) were added to the wells.

DC and T cells were co-cultured at 37ºC for 72 h, then cells were harvested and stained for flow cytometry.

### Phagocytosis assay

Cells were processed as for the DC-T cell assay, but the MOI was reduced to 10 and cells were plated after the gentamycin incubation for CFU enumeration.

### Tissue Processing

#### Lymph nodes and Lymphoid cells

Lymph nodes (skin-draining and colon-draining) were harvested and then processed over sterile wire mesh in 2 mL of complete RPMI media before cell isolation and tetramer staining in PBS.

#### Lymph nodes and myeloid populations

Harvested lymph nodes were incubated for 20 minutes at 37ºC in RPMI with one mL of digestion solution (2 mg/mL collagenase I, 2 mg/mL collagenase IV, 0.1 mg/mL DNase in RPMI with 1%HEPES,1%penicillin-streptomycin and10% fetal calf serum), resuspended, and further incubated for 15 minutes at 37ºC. Cell suspension was quenched in 9 mL of complete medium and filtered through a 40 μm sterile cell strainer before counting and staining.

#### Mouse skin processing

Back skin was harvested, lightly defatted, and then washed in PBS by extensive vortexing to reduce free floating bacteria. Skin was then dried and minced with scissors to a fine consistency before tissue digestion in 3 mL complete RPMI (RPMI plus 10% fetal calf serum, 1% penicillin-streptomycin, β-mercaptoethanol, glutamate, sodium pyruvate, HEPES and non-essential amino acids) then supplemented with 2 mg/mL collagenase XI, 0.5 mg/mL hyaluronidase, 0.1 mg/mL DNase I. When looking at bacterial uptake in vivo, cytochalasin D (Cayman Chemicals, # 11330) at 2.5 μg/mL was added. Digested skin samples were then incubated, with shaking, at 37ºC for 45 minutes before quenching with 15 mL of complete RPMI media and shaking by hand for 30 seconds. Skin cell suspensions were filtered through sterile cell strainers (100 μm followed by 40 μm).

#### Epidermal preparations

Washed and defatted back skin were floated on a solution of Trypsin 0.5% + 2.5 μg/mL of cytochalasin D for 45 minutes at 37ºC. Tissues were transferred to 1 mL of complete RPMI and the epithelial cells were scratched off the tissue. 9 mL of complete RPMI was added and cells filtered through a 40 μm sterile cell strainer.

#### Human Skin Processing

Skin tissued were finely minced with scissors and incubated in a digestion cocktail containing collagenase IV (3.2 mg.ml^−1^, Worthington, LS004186) and DNAse (20 μg ml^−1^, MilliporeSigma, DN251G) diluted in RPMI with 10% fetal bovine serum (FBS), 1% HEPES, 1% non-essential amino acids, 1% Glutamax and 1% penicillin–streptomycin with cytochalasin D 2.5 μg/mL for 2 h at 37 ºC. Digests were briefly shaken and passed through a 100-µm strainer to yield a single-cell suspension.

#### Cell counting

All tissues were re-suspended in 1 mL PBS and 25 μL of cell suspension was mixed with 25 μL of AccuCheck counting beads (Invitrogen,Catalog No. PCB100) for calculating absolute numbers of cells.

### Flow Cytometry

#### Antibody Staining of Myeloid Populations

Cells were stained in PBS for 20 minutes at 4ºC with a Live/Dead marker (Ghost Dye Violet510, Tonbo Biosciences, Catalog No. 13-0870-T100). Surface antibodies were added in blocking solution for 30 minutes at 4ºC. Cells were washed and fixed in Paraformaldehyde (PFA) for 30 minutes at 4ºC.

#### Antibody Staining of Lymphoid Populations

Cells were stained in blocking solution with a Live/Dead marker (Ghost Dye Violet510, Tonbo Biosciences) for 30 minutes at 4ºC. For intracellular staining, cells were fixed and permeabilized using the Foxp3 staining kit (eBioscience, Catalog No. 00-5523-00) buffer for 30 min. at 4ºC then stained in permeabilization buffer for 30 minutes at 4ºC.

#### Cell Re-stimulation for Cytokines

After isolation, skin cells were stimulated with Tonbo kit (1/100 in 1 mL) for 3-6 hours before being processed for staining. A subset of cells was incubated in Brefeldin A to be used as an unstimulated control for gating.

#### Spectral Flow Cytometry

Cells were incubated with a live-dead dye (ViaDye Red, Cytek Bio) for 20 minutes at 25ºC. CCR7 antibody was added for 15 minutes at 37 ºC. Surface staining in blocking solution was then added for 30 minutes at 4ºC. Cells were fixed with 4% PFA for 15 minutes at 25ºC.

#### BandPath Flow cytometry

Stained cells were run on a Fortessa (BD Biosciences) or an Aurora (Cytek) in the UCSF Flow Cytometry Core. Flow cytometry data was analyzed using FlowJo software (FlowJo, LLC).

#### Tetramer Staining and Enrichment

To identify 2w-specific cells, cell suspensions were pelleted and then stained for 1 hour at room temperature (15–25ºC), while protected from light, with a 2W1S:I-Ab–streptavidin-phycoerythrin (PE) tetramer at a concentration of 10 nM. Skin was directly stained for other surface and intracellular markers as described above. For LN samples, the tetramer-bound fraction was enriched via an adapted protocol of the EasySep PE Selection Kit II (StemCell Technologies, Inc.) developed by Marc Jenkins’ lab. In brief, 6.25 μL of EasySep PE selection cocktail was added to each sample in a total volume of 500 mL and then incubated, while protected from light, at room temperature for 15 minutes. Cells were incubated for an additional 10 minutes after addition of 6.25 μL of EasySep magnetic particles. Finally, cell suspensions were brought up to a total volume of 2.5 mL with PBS and placed into the EasySep magnet for 5 minutes at room temperature. Supernatants (unbound fractions) were poured off into another collection tube. The positively-selected cells (bound fraction) and unbound fraction for each sample were taken for cell counting and staining.

### Human Foreskin Explant

#### Explant methodology

Fresh foreskins from elective circumcision of newborn male infants were obtained as deidentified samples, washed in RPMI and cut to generate two explants. Silicone rings were glued to the skin explant using Kwik-Sil silicone glue. Explants were floated in a 12 mm Milicell (Sigma) hanging well on a neutralized solution of 2 mg/mL collagen in RPMI (Thermo Scientific) at 37ºC for 30 min until the explant was firmly embedded in a collagen hydrogel. It was then placed in a 12 well plate with the lower chamber filled with RPMI. 5 mL of a 24h culture of *S*.*epi-*zsgreen was washed and resuspended in 10 mL of RPMI. 50 µL of this bacterial solution was added to the center of the silicone ring. Bacteria were left to interact with the tissue for 30 min at 37ºC, and the solution of bacteria was then removed to restore an air-skin interface. The explant was left for an additional 4 h at 37ºC until tissue processing as above.

### Mouse scRNAseq

Live singlet dendritic cells (CD45^+^ MHCII^+^ CD11c^+^) loaded (zsgreen^+^) or not (zsgreen^neg^) from digested back skin or skin draining lymph nodes of control or *S-epi-*zsgreen colonized D10 neonatal mice were run on a BD FACSAriaII using a 100-µm nozzle and sorted into RPMI medium supplemented with 10% FBS. Cells were sorted from 7-10 pooled mice and different conditions were run on separate lanes of a 10X Chromium chip with 3′ v.2 chemistry (10X Genomics) as per the manufacturer’s instructions by the UCSF Institute for Human Genetics Sequencing Core. Libraries were sequenced on an Illumina Novaseq 6000. Fastq files were aligned to the mm10 reference genome, and barcode matrices were generated using CellRanger 2.1. For Cite-seq, Totalseq-A antibodies were added at the same time antibodies for the sort were added. The cite-seq library was generated according to the manufacturer protocol and sequenced independently.

Downstream data analysis, including clustering, visualizations and exploratory analyses, were performed using Seurat 4.0. Cells with <200 features or more than 5500, or >6% mitochondrial genes were filtered out during preprocessing. Samples sequenced in parallel lanes were merged together for downstream analysis. Principal component analysis (PCA) and uniform manifold approximation and projection (UMAP) were run on the RNA data, and an initial low-resolution clustering was generated using the first 25 principal components. Datasets of figure 1 were clustered based on the RNA transcripts but the protein expression obtained with Cite-seq was used to identify DC clusters using the same defining markers delineated in the flow cytometry gating strategy shown in Figure S3. Datasets of Fig. 5 used RNA signature based on classical defining markers (*Xcr1* for DC1, *Epcam* Langerhans cells and *Irf4* together with *Itgam* for CD11b^hi^ DC2 and CD11b^lo^ DC2, *Ccr7* for CCR7+ DCs).

### Human scRNAseq

Live myeloid cells (CD45^+^ MHCII^+^ CD11c^+^) loaded (zsgreen^+^) or not (zsgreen^neg^) from a baby foreskin explant colonized with *S*.*epi*-zsgreen were sorted on a BD FACS AriaII using a 100-µm nozzle and sorted into RPMI medium supplemented with 10% FBS. Cells were sorted from 8 explants corresponding to duplicated explants from 4 independent fresh donors (<24h). Downstream processing followed the above mouse scRNAseq protocol, except for the quality control thresholds. Cells below 200 features and above 5000 features or with a percentage of mitochondrial genes above 25% were excluded. Genes used to delineate clusters are shown in Fig. S5. For Fig. 5F, the dendritic cell cluster of Fig. 5E was re-clustered to identify sub-population of dendritic cells.

### Quantitative RT-PCR

Indicated populations were sorted into RLT Plus lysis buffer (Qiagen) and stored at -80°C, then processed using Allprep DNA/RNA micro kit (Qiagen) per manufacturer’s protocol. For qPCR analyses, RNA was reverse transcribed using SuperScript III cDNA synthesis kit (ThermoFisher) and amplified using Power SYBR Green PCR master mix (ThermoFisher). *Aldh1a2* primers: 5′-GACTTGTAGCAGCTGTCTTCACT-3′, forward, and 5′-TCACCCATTTCTCTCCCATTTCC-3′, reverse. *Rsp17* gene was used as a housekeeping gene and amplified with the following primers: ATTGAGGTGGATCCCGACAC, forward; TGCCAACTGTAGGCTGAGTG, reverse.

### AldeRed Assay

10,000-100,000 sorted cells were plated in a 96 well plates. Staining with AldeRed ALDH detection assay (Sigma) was done following manufacturer’s recommendation with minor modifications. To account for the small number of cells, volumes were decreased from the standard protocol recommendations to 100 µL and the reagents AldeRed and DEAB were added at 1 µL per well. Cells were incubated for an hour at 37ºC.

### Data availability and software used

Schematic were made with Biorender (https://biorender.com). The volcano plot was generated with VolcaNoseR (Goedhart and Luijsterburg, 2020). The panther analysis (http://www.pantherdb.org/) was used with statistical overrepresentation to “GO biological process complete”. GraphPad (Prism) 9.3.1 was used. Rstudio v4 was used for scRNAseq analysis.

## References

Akagbosu, B., Tayyebi, Z., Shibu, G., Iza, Y.A.P., Deep, D., Parisotto, Y.F., Pasolli, H.A., Elmentaite, R., Knott, M., Hemmers, S., et al. (2022). A novel lineage of RORγt+Aire+ antigen presenting cells promotes peripheral generation of intestinal regulatory T cells and tolerance during early life. BioRxiv 2022.02.26.481148. https://doi.org/10.1101/2022.02.26.481148.

Allgaier, H., Jung, G., Werner, R.G., Schneider, U., and Zähner, H. (1986). Epidermin: sequencing of a heterodetic tetracyclic 21-peptide amide antibiotic. European Journal of Biochemistry / FEBS 160, 9–22..

Augustin, J., and Gotz, F. (1990). Transformation of Staphylococcus epidermidis and other staphylococcal species with plasmid DNA by electroporation. FEMS Microbiology Letters 54, 203–207..

Bar-On, L., Birnberg, T., Kim, K., and Jung, S. (2011). Dendritic cell-restricted CD80/86 deficiency results in peripheral regulatory T-cell reduction but is not associated with lymphocyte hyperactivation. European Journal of Immunology 41, 291–298. https://doi.org/10.1002/EJI.201041169.

Benson, M.J., Pino-Lagos, K., Rosemblatt, M., and Noelle, R.J. (2007). All-trans retinoic acid mediates enhanced T reg cell growth, differentiation, and gut homing in the face of high levels of co-stimulation. Journal of Experimental Medicine 204, 1765–1774. https://doi.org/10.1084/JEM.20070719.

Bobr, A., Olvera-Gomez, I., Igyarto, B.Z., Haley, K.M., Hogquist, K.A., and Kaplan, D.H. (2010). Acute ablation of Langerhans cells enhances skin immune responses. Journal of Immunology (Baltimore, Md.: 1950) 185, 4724–4728. https://doi.org/10.4049/jimmunol.1001802.

Breed, E.R., Vobořil, M., Ashby, K.M., Martinez, R.J., Qian, L., Wang, H., Salgado, O.C., O’Connor, C.H., and Hogquist, K.A. (2022). Type 2 cytokines in the thymus activate Sirpα+ dendritic cells to promote clonal deletion. Nature Immunology 2022 1–10. https://doi.org/10.1038/s41590-022-01218-x.

Brodin, P. (2022). Immune-microbe interactions early in life: A determinant of health and disease long term. Science (1979) 376, 945–950. https://doi.org/10.1126/SCIENCE.ABK2189.

Brough, H.A., Lanser, B.J., Sindher, S.B., Teng, J.M.C., Leung, D.Y.M., Venter, C., Chan, S.M., Santos, A.F., Bahnson, H.T., Guttman-Yassky, E., et al. (2022). Early intervention and prevention of allergic diseases. Allergy 77, 416–441. https://doi.org/10.1111/ALL.15006.

Brown, C.C., Gudjonson, H., Pritykin, Y., Deep, D., Lavallée, V.-P., Mendoza, A., Fromme, R., Mazutis, L., Ariyan, C., Leslie, C., et al. (2019a). Transcriptional Basis of Mouse and Human Dendritic Cell Heterogeneity. Cell 179, 846-863.e24.

Brown, C.C., Gudjonson, H., Pritykin, Y., Deep, D., Lavallée, V.-P., Mendoza, A., Fromme, R., Mazutis, L., Ariyan, C., Leslie, C., et al. (2019b). Transcriptional Basis of Mouse and Human Dendritic Cell Heterogeneity. Cell 179, 846-863.e24.

Fu, H., Song, S., Liu, F., Ni, Z., Tang, Y., Shen, X., Xiao, L., Ding, G., and Wang, Q. (2009). Dendritic cells transduced with SOCS1 gene exhibit regulatory DC properties and prolong allograft survival. Cell. Mol. Immunol. 6, 87–95.

Goedhart, J., and Luijsterburg, M.S. (2020). VolcaNoseR is a web app for creating, exploring, labeling and sharing volcano plots. Sci. Rep. 10, 20560.

Mouse and Human Dendritic Cell Heterogeneity. Cell 179, 846-863.e24. https://doi.org/10.1016/j.cell.2019.09.035.

Byrd, A.L., Belkaid, Y., and Segre, J.A. (2018). The human skin microbiome. Nature Reviews Microbiology 16, 143–155. https://doi.org/10.1038/nrmicro.2017.157.

Chaudhry, A., Samstein, R.M., Treuting, P., Liang, Y., Pils, M.C., Heinrich, J.-M., Jack, R.S., Wunderlich, F.T., Brüning, J.C., Müller, W., et al. (2011). Interleukin-10 signaling in regulatory T cells is required for suppression of Th17 cell-mediated inflammation. Immunity 34, 566. https://doi.org/10.1016/J.IMMUNI.2011.03.018.

Coombes, J.L., Siddiqui, K.R.R., Arancibia-Carcamo, C. v, Hall, J., Sun, C.M., Belkaid, Y., and Powrie, F. (2007). A functionally specialized population of mucosal CD103+ DCs induces Foxp3+ regulatory T cells via a TGF- and retinoic acid dependent mechanism. J Exp Med 204, 1757–1764. https://doi.org/10.1093/nar/29.9.e45.

Domogalla, M.P., Rostan, P. v, Raker, V.K., and Steinbrink, K. (2017). Tolerance through Education: How Tolerogenic Dendritic Cells Shape Immunity. Frontiers in Immunology 8. https://doi.org/10.3389/fimmu.2017.01764.

Francisco, L.M., Salinas, V.H., Brown, K.E., Vanguri, V.K., Freeman, G.J., Kuchroo, V.K., and Sharpe, A.H. (2009). PD-L1 regulates the development, maintenance, and function of induced regulatory T cells. Journal of Experimental Medicine 206, 3015–3029. https://doi.org/10.1084/JEM.20090847.

Gallo, R.L. (2017). Human Skin Is the Largest Epithelial Surface for Interaction with Microbes. The Journal of Investigative Dermatology 137, 1213–1214. https://doi.org/10.1016/j.jid.2016.11.045.

Guilliams, M., Crozat, K., Henri, S., Tamoutounour, S., Grenot, P., Devilard, E., de Bovis, B., Alexopoulou, L., Dalod, M., and Malissen, B. (2010). Skin-draining lymph nodes contain dermis-derived CD103− dendritic cells that constitutively produce retinoic acid and induce Foxp3+ regulatory T cells. Blood 115, 1958–1968. https://doi.org/10.1182/blood-2009-09-245274.

Haniffa, M., Gunawan, M., and Jardine, L. (2015). Human skin dendritic cells in health and disease. Journal of Dermatological Science 77, 85. https://doi.org/10.1016/J.JDERMSCI.2014.08.012.

Harris-Tryon, T.A., and Grice, E.A. (2022). Microbiota and maintenance of skin barrier function. Science (1979) 376, 940–945. https://doi.org/10.1126/SCIENCE.ABO0693.

Ji, S.J., Zhuang, B.Q., Falco, C., Schneider, A., Schuster-Gossler, K., Gossler, A., and Sockanathan, S. (2006). Mesodermal and neuronal retinoids regulate the induction and maintenance of limb innervating spinal motor neurons. Developmental Biology 297, 249–261. https://doi.org/10.1016/J.YDBIO.2006.05.015.

Kim, T.G., Kim, S.H., Park, J., Choi, W., Sohn, M., Na, H.Y., Lee, M., Lee, J.W., Kim, S.M., Kim, D.Y., et al. (2018). Skin-Specific CD301b+ Dermal Dendritic Cells Drive IL-17−Mediated Psoriasis-Like Immune Response in Mice. Journal of Investigative Dermatology 138, 844–853. https://doi.org/10.1016/J.JID.2017.11.003.

Knoop, K.A., Gustafsson, J.K., McDonald, K.G., Kulkarni, D.H., Coughlin, P.E., McCrate, S., Kim, D., Hsieh, C.S., Hogan, S.P., Elson, C.O., et al. (2017). Microbial antigen encounter during a preweaning interval is critical for tolerance to gut bacteria. Science Immunology 2. https://doi.org/10.1126/SCIIMMUNOL.AAO1314.

Kulkarni, D.H., Gustafsson, J.K., Knoop, K.A., McDonald, K.G., Bidani, S.S., Davis, J.E., Floyd, A.N., Hogan, S.P., Hsieh, C.-S., and Newberry, R.D. (2019). Goblet cell associated antigen passages support the induction and maintenance of oral tolerance. Mucosal Immunology 2019 13:2 13, 271–282. https://doi.org/10.1038/s41385-019-0240-7.

Kumamoto, Y., Linehan, M., Weinstein, J.S., Laidlaw, B.J., Craft, J.E., and Iwasaki, A. (2013). CD301b^+^ dermal dendritic cells drive T helper 2 cell-mediated immunity. Immunity 39, 733–743. https://doi.org/10.1016/j.immuni.2013.08.029.

Larange, A., and Cheroutre, H. (2016). Retinoic Acid and Retinoic Acid Receptors as Pleiotropic Modulators of the Immune System. Annu Rev Immunol 34, 369–394. https://doi.org/10.1146/ANNUREV-IMMUNOL-041015-055427.

Leech, J.M., Dhariwala, M.O., Lowe, M.M., Chu, K., Merana, G.R., Cornuot, C., Weckel, A., Ma, J.M., Leitner, E.G., Gonzalez, J.R., et al. (2019). Toxin-Triggered Interleukin-1 Receptor Signaling Enables Early-Life Discrimination of Pathogenic versus Commensal Skin Bacteria. Cell Host and Microbe 26, 795–809.e5. https://doi.org/10.1016/j.chom.2019.10.007.

Linehan, J.L., Dileepan, T., Kashem, S.W., Kaplan, D.H., Cleary, P., and Jenkins, M.K. (2015). Generation of Th17 cells in response to intranasal infection requires TGF-β1 from dendritic cells and IL-6 from CD301b+ dendritic cells. Proc Natl Acad Sci U S A 112, 12782–12787. https://doi.org/10.1073/PNAS.1513532112.

Loschko, J., Rieke, G.J., Schreiber, H.A., Meredith, M.M., Yao, K.H., Guermonprez, P., and Nussenzweig, M.C. (2016). Inducible targeting of cDCs and their subsets in vivo. Journal of Immunological Methods 434, 32–38. https://doi.org/10.1016/J.JIM.2016.04.004.

Maier, B., Leader, A.M., Chen, S.T., Tung, N., Chang, C., LeBerichel, J., Chudnovskiy, A., Maskey, S., Walker, L., Finnigan, J.P., et al. (2020). A conserved dendritic-cell regulatory program limits antitumour immunity. Nature 580, 257–262. https://doi.org/10.1038/s41586-020-2134-y.

Matz, M. v., Fradkov, A.F., Labas, Y.A., Savitsky, A.P., Zaraisky, A.G., Markelov, M.L., and Lukyanov, S.A. (1999). Fluorescent proteins from nonbioluminescent Anthozoa species. Nature Biotechnology 1999 17:10 17, 969–973. https://doi.org/10.1038/13657.

Mayer, J.U., Hilligan, K.L., Chandler, J.S., Eccles, D.A., Old, S.I., Domingues, R.G., Yang, J., Webb, G.R., Munoz-Erazo, L., Hyde, E.J., et al. (2021). Homeostatic IL-13 in healthy skin directs dendritic cell differentiation to promote TH2 and inhibit TH17 cell polarization. Nature Immunology 2021 22:12 22, 1538–1550. https://doi.org/10.1038/s41590-021-01067-0.

McGovern, N., Shin, A., Low, G., Low, D., Duan, K., Yao, L.J., Msallam, R., Low, I., Shadan, N.B., Sumatoh, H.R., et al. (2017). Human fetal dendritic cells promote prenatal T-cell immune suppression through arginase-2. Nature 546, 662–666. https://doi.org/10.1038/nature22795.

Merana, G.R., Dwyer, L.R., Dhariwala, M.O., Weckel, A., Gonzalez, J.R., Okoro, J.N., Cohen, J.N., Tamaki, C.M., Han, J., Tasoff, P., et al. (2022). Intestinal inflammation alters the antigen-specific immune response to a skin commensal. Cell Reports 39, 110891. https://doi.org/10.1016/J.CELREP.2022.110891.

Mucida, D., Park, Y., Kim, G., Turovskaya, O., Scott, I., Kronenberg, M., and Cheroutre, H. (2007). Reciprocal TH17 and Regulatory T Cell Differentiation Mediated by Retinoic Acid. Science (1979) 317, 256–260. https://doi.org/10.1126/SCIENCE.1145697.

Mucida, D., Pino-Lagos, K., Kim, G., Nowak, E., Benson, M.J., Kronenberg, M., Noelle, R.J., and Cheroutre, H. (2009). Retinoic Acid Can Directly Promote TGF-β-Mediated Foxp3+ Treg Cell Conversion of Naive T Cells. Immunity 30, 471–472. https://doi.org/10.1016/J.IMMUNI.2009.03.008.

Nirschl, C.J., Suárez-Fariñas, M., Izar, B., Prakadan, S., Dannenfelser, R., Tirosh, I., Liu, Y., Zhu, Q., Devi, K.S.P., Carroll, S.L., et al. (2017). IFNγ-Dependent Tissue-Immune Homeostasis Is Co-opted in the Tumor Microenvironment. Cell 170, 127–141.e15. https://doi.org/10.1016/j.cell.2017.06.016.

Qian, C., Qian, L., Yu, Y., An, H., Guo, Z., Han, Y., Chen, Y., Bai, Y., Wang, Q., and Cao, X. (2013). Fas Signal Promotes the Immunosuppressive Function of Regulatory Dendritic Cells via the ERK/β-Catenin Pathway *. Journal of Biological Chemistry 288, 27825–27835. https://doi.org/10.1074/JBC.M112.425751.

Rios, D., Wood, M.B., Li, J., Chassaing, B., Gewirtz, A.T., and Williams, I.R. (2015). Antigen sampling by intestinal M cells is the principal pathway initiating mucosal IgA production to commensal enteric bacteria. Mucosal Immunology 2016 9:4 9, 907–916. https://doi.org/10.1038/mi.2015.121.

Salomon, B., Lenschow, D.J., Rhee, L., Ashourian, N., Singh, B., Sharpe, A., and Bluestone, J.A. (2000). B7/CD28 Costimulation Is Essential for the Homeostasis of the CD4+CD25+ Immunoregulatory T Cells that Control Autoimmune Diabetes. Immunity 12, 431–440. https://doi.org/10.1016/S1074-7613(00)80195-8.

Scharschmidt, T.C., Vasquez, K.S., Truong, H.-A., Gearty, S. v, Pauli, M.L., Nosbaum, A., Gratz, I.K., Otto, M., Moon, J.J., Liese, J., et al. (2015). A wave of regulatory T cells into neonatal skin mediates tolerance to commensal microbes. Immunity 43..

Shin, H., Kumamoto, Y., Gopinath, S., and Iwasaki, A. (2016). CD301b+ dendritic cells stimulate tissue-resident memory CD8+ T cells to protect against genital HSV-2. Nature Communications 7, 1–10. https://doi.org/10.1038/ncomms13346.

Steinman, R.M., Hawiger, D., and Nussenzweig, M.C. (2003). Tolerogenic Dendritic Cells*. https://Doi.Org/10.1146/Annurev.Immunol.21.120601.141040 21, 685–711. https://doi.org/10.1146/ANNUREV.IMMUNOL.21.120601.141040.

Sun, C.-M.M., Hall, J.A., Blank, R.B., Bouladoux, N., Oukka, M., Mora, J.R., and Belkaid, Y. (2007). Small intestine lamina propria dendritic cells promote de novo generation of Foxp3 T reg cells via retinoic acid. Journal of Experimental Medicine 204, 1775–1785. https://doi.org/10.1084/jem.20070602.

Tatsumi, N., Codrington, A.L., El-Fenej, J., Phondge, V., and Kumamoto, Y. (2021). Effective CD4 T cell priming requires repertoire scanning by CD301b+ migratory cDC2 cells upon lymph node entry. Science Immunology 6. https://doi.org/10.1126/sciimmunol.abg0336.

Villablanca, E.J., Wang, S., Calisto, J. de, Gomes, D.C.O., Kane, M.A., Napoli, J.L., Blaner, W.S., Kagechika, H., Blomhoff, R., Rosemblatt, M., et al. (2011). MyD88 and Retinoic Acid Signaling Pathways Interact to Modulate Gastrointestinal Activities of Dendritic Cells. Gastroenterology 141, 176–185. https://doi.org/10.1053/J.GASTRO.2011.04.010.

Vogel, A., Martin, K., Soukup, K., Halfmann, A., Kerndl, M., Brunner, J.S., Hofmann, M., Oberbichler, L., Korosec, A., Kuttke, M., et al. (2022). JAK1 signaling in dendritic cells promotes peripheral tolerance in autoimmunity through PD-L1-mediated regulatory T cell induction. Cell Reports 38, 110420. https://doi.org/10.1016/J.CELREP.2022.110420.

Wang, J., Lareau, C.A., Bautista, J.L., Gupta, A.R., Sandor, K., Germino, J., Yin, Y., Arvedson, M.P., Reeder, G.C., Cramer, N.T., et al. (2021). Single-cell multiomics defines tolerogenic extrathymic Aire-expressing populations with unique homology to thymic epithelium. Science Immunology 6. https://doi.org/10.1126/SCIIMMUNOL.ABL5053.

Wang, S., Villablanca, E.J., Calisto, J. de, Gomes, D.C.O., Nguyen, D.D., Mizoguchi, E., Kagan, J.C., Reinecker, H.-C., Hacohen, N., Nagler, C., et al. (2011). MyD88-Dependent TLR1/2 Signals Educate Dendritic Cells with Gut-Specific Imprinting Properties. The Journal of Immunology 187, 141–150. https://doi.org/10.4049/JIMMUNOL.1003740.

Xiao, S., Jin, H., Korn, T., Liu, S.M., Oukka, M., Lim, B., and Kuchroo, V.K. (2008). Retinoic Acid Increases Foxp3+ Regulatory T Cells and Inhibits Development of Th17 Cells by Enhancing TGF-β-Driven Smad3 Signaling and Inhibiting IL-6 and IL-23 Receptor Expression. The Journal of Immunology 181, 2277–2284. https://doi.org/10.4049/JIMMUNOL.181.4.2277.

Yamazaki, C., Sugiyama, M., Ohta, T., Hemmi, H., Hamada, E., Sasaki, I., Fukuda, Y., Yano, T., Nobuoka, M., Hirashima, T., et al. (2013a). Critical Roles of a Dendritic Cell Subset Expressing a Chemokine Receptor, XCR1. The Journal of Immunology 190, 6071–6082. https://doi.org/10.4049/JIMMUNOL.1202798.

Yamazaki, C., Sugiyama, M., Ohta, T., Hemmi, H., Hamada, E., Sasaki, I., Fukuda, Y., Yano, T., Nobuoka, M., Hirashima, T., et al. (2013b). Critical Roles of a Dendritic Cell Subset Expressing a Chemokine Receptor, XCR1. The Journal of Immunology 190, 6071–6082. https://doi.org/10.4049/JIMMUNOL.1202798.

Zegarra-Ruiz, D.F., Kim, D. v., Norwood, K., Kim, M., Wu, W.-J.H., Saldana-Morales, F.B., Hill, A.A., Majumdar, S., Orozco, S., Bell, R., et al. (2021). Thymic development of gut-microbiota-specific T cells. Nature 2021 594:7863 594, 413–417. https://doi.org/10.1038/s41586-021-03531-1.

