## Supplementary figures for "Early life tolerance depends on a subset of specialized dendritic cells and is reinforced by the skin microbiota"

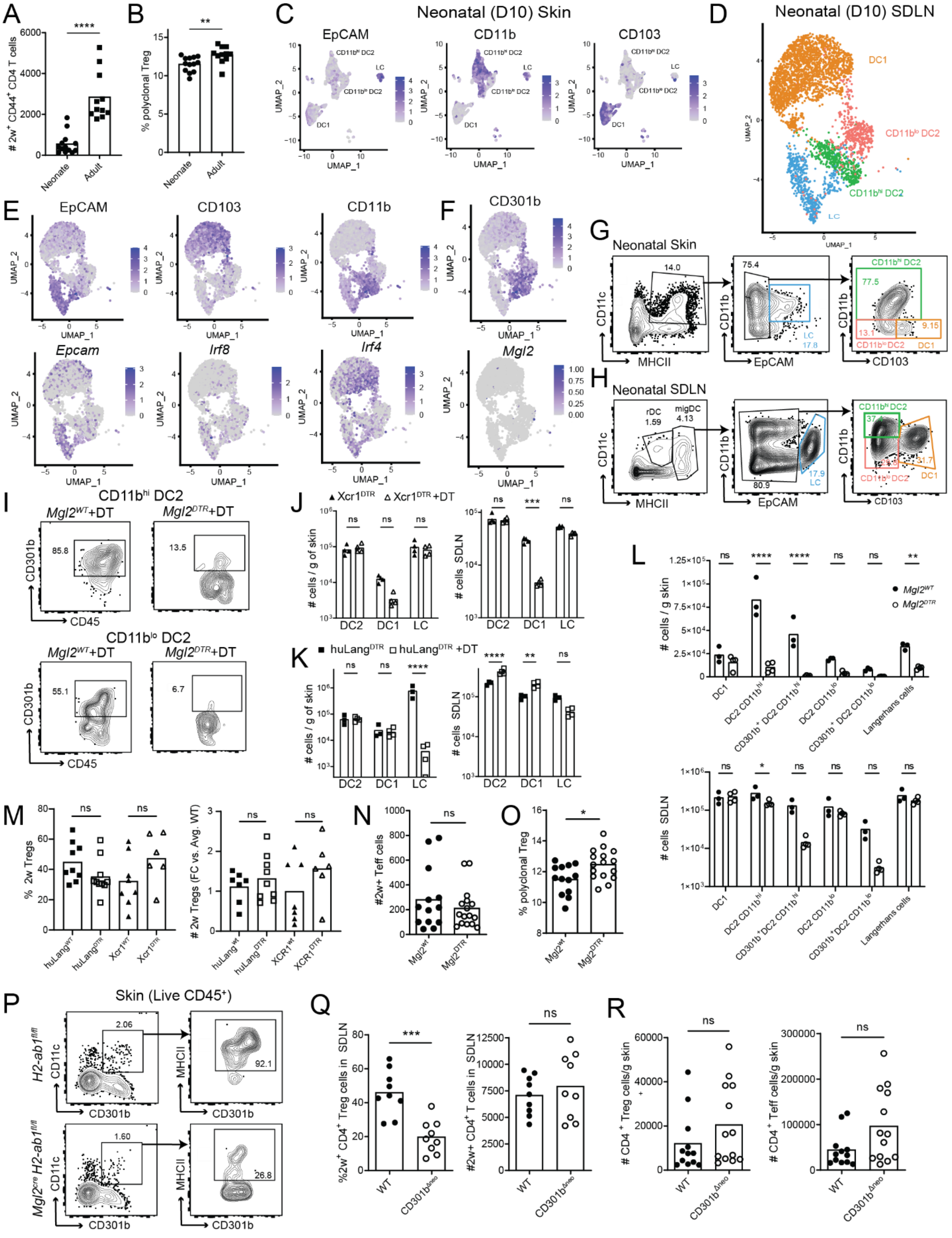

**Figure S1: Related to main Figure 1** (A) Total number of 2w$^{+}$CD4$^{+}$CD44$^{+}$ and (B) percentage of polyclonal CD4$^{+}$ FoxP3$^{+}$ cells in the SDLN of adult and neonates after primary *S.* epi-2w challenge. (C) Protein level expression of DC markers in D10 skin by CITE-Seq. (D) UMAP of major DC populations among MHCII$^{hi}$CD11c$^{low\text{-}high}$ migratory DC (migDC) sorted from SDLN of neonatal (D10) pups and submitted for CITE-Seq. (E) RNA- (bottom) and protein- (top) level expression of major surface markers and of (F) *Mgl2*/CD301b among D10 SDLN migDC. (G-H) Gating strategy to quantify major skin DC populations by flow cytometry in D10 (G) skin and (H) SDLN (pre-gated on live CD45+ cells). Color code correspond to UMAP in Figs. 1C and S1D. (I-L) Representative plots in the skin and (I) quantification of DC populations 24h after DT injection in D11 mice of the indicated genotypes in the skin and SDLN. One of two representative experiments is shown. (M-O) Refers to Fig. 1H-L. (M) Percentage of Tregs among SDLN 2w$^{+}$CD4$^{+}$CD44$^{+}$ (left) and the fold change in number of these 2w+ Tregs (right) relative to the average of controls in the indicated genotypes. (N) Total number of 2w$^{+}$CD4$^{+}$CD44$^{+}$ FoxP3$^{neg}$ cells in the SDLN. (O) Percentage of FoxP3$^{+}$ cells among polyclonal CD4$^{+}$T cells in the SLDN. (P) Sample gating to of D21 skin showing efficiency of MHCII deletion among CD301b$^{+}$ cells in *Mgl2*$^{Cre}$*H2-ab1*$^{fl/fl}$ mice. Mann-Whitney test except in J-L, Two-Way ANOVA. *p < 0.05, **p < 0.01, ***p < 0.001, ****p<0.0001, ns = not significant. FC vs. Avg. WT= fold change to the average of the WT.

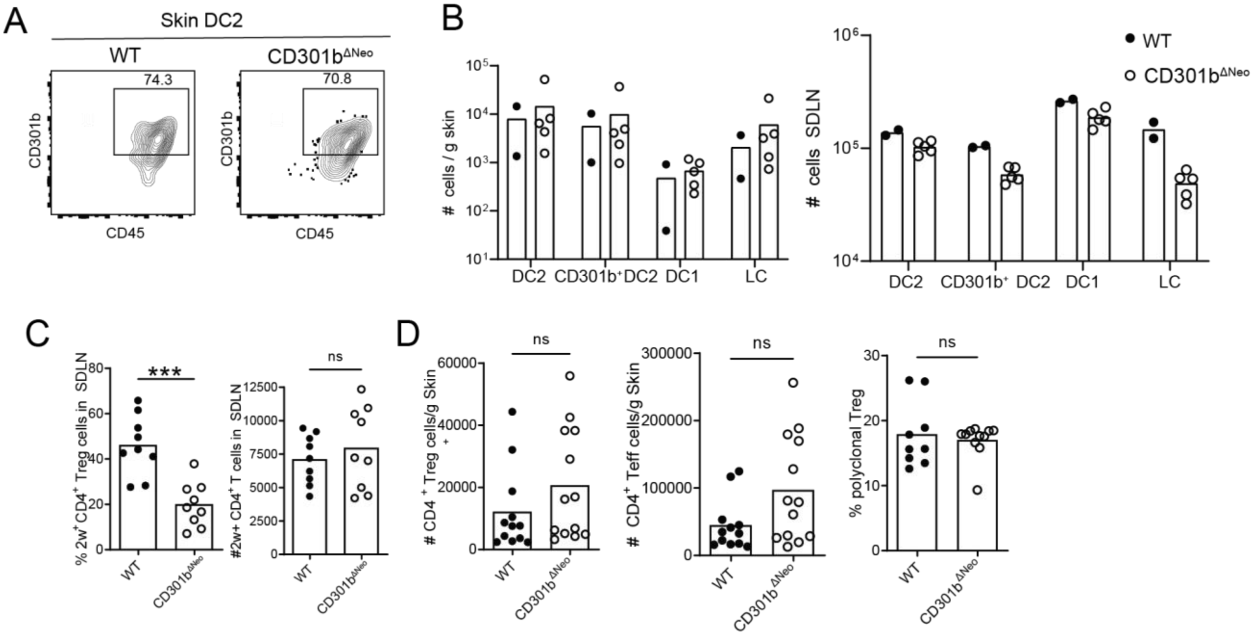


**Figure S2: Related to main Figure 2 (A-B)** Two weeks after neonatal depletion of $CD301b^+$ cells, skin and SDLN were harvested. (A) Representative percentage of $CD301b^+$ DC2 in WT vs neonatally depleted mice ($CD301b^{\Delta Neo}$) and (B) quantification in the skin (left) and SDLN (right) of the DC populations. (C-D) refers to Fig. 2A-E. (C) Percentage (left) and total number (right) of $2w^+CD4^+CD44^+$ $FoxP3^+$ in SDLN. (D) Total number (left) and percentage (right) of polyclonal $CD4^+$ $FoxP3^+$ (left) and $CD4^+$ $FoxP3^{neg}$ per gram of skin (center). Dots represent individual mice. Mann-Whitney test, *$p < 0.05$, **$p < 0.01$, ***$p < 0.001$, ns = not significant.

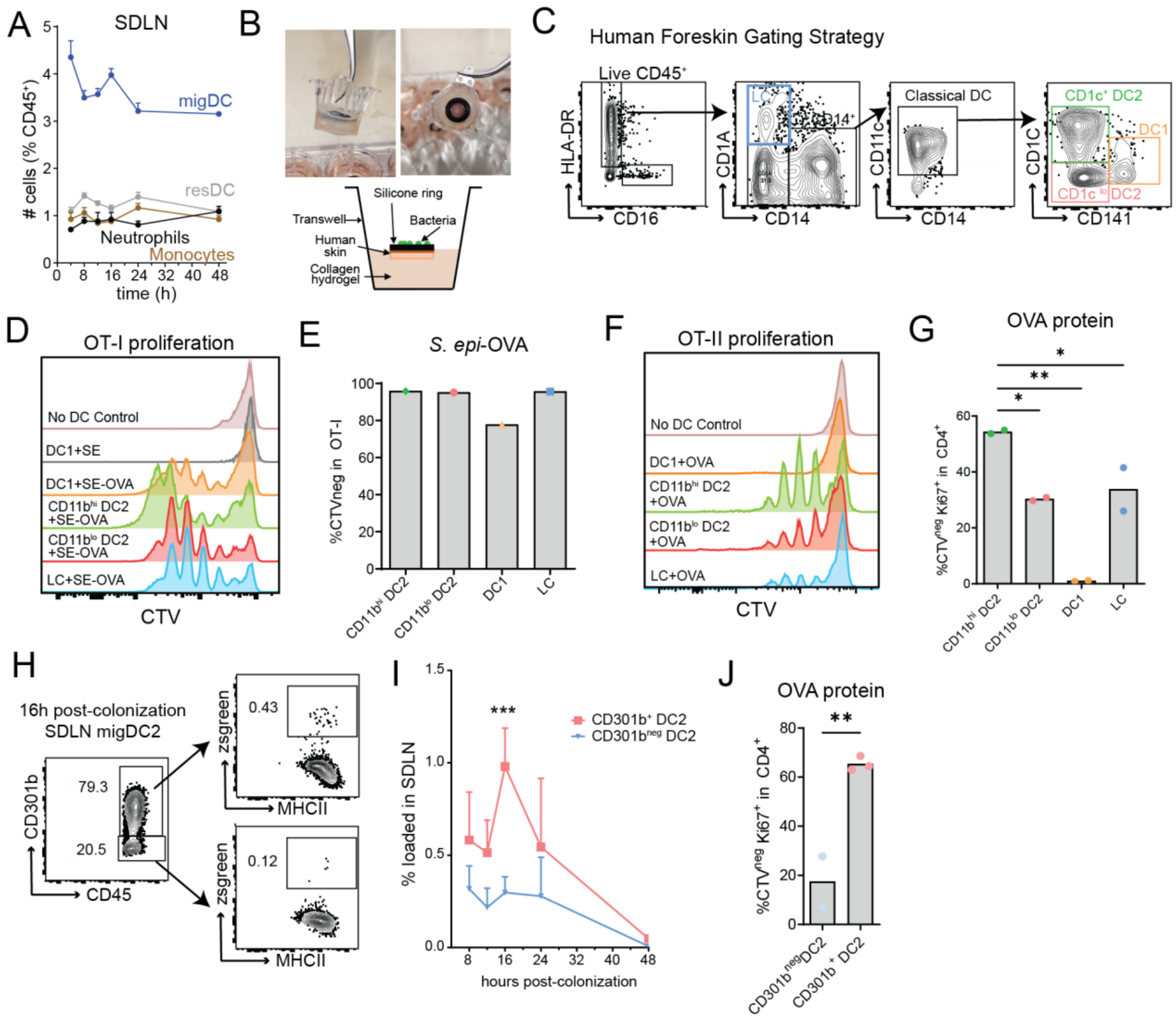


**Figure S3: Related to main Figure 3** (A) Counts of major myeloid populations in the SDLN in the hours after skin colonization by *S. epidermidis*. (B) Schematic and pictures of explant system used to colonize human neonatal foreskin. In brief, a silicone ring is sealed to an 8mm punch biopsy of freshly collected tissue and the explant is embedded in collagen hydrogel within a hanging transwell. *S.epi*-zsgreen is added in the center of the ring for 4h at 37C during which bacteria only access tissue from the epidermal surface. (C) Gating strategy for major DC populations in human foreskin (pre-gated on Live CD45$^+$). (D-G, J) Same experimental set up as in Fig. 3H but sorted SDLN DCs were incubated with OVA protein prior to OT-I (D-E) or OT-II (F-G and J) co-culture. (D and F) Histogram and (E, G and J) quantification of T cell proliferation, as measured by CTV dilution. (H) Sample gating and (I) quantification of bacterial uptake by CD301b$^+$ vs CD301b$^{neg}$ DC2 in the SDLN (pre-gated as in Fig. S1H inclusive of CD11b hi and lo subsets. Two-way ANOVA with Šídák's multiple comparisons test. **p < 0.01, ****p<0.0001, ns = not significant.

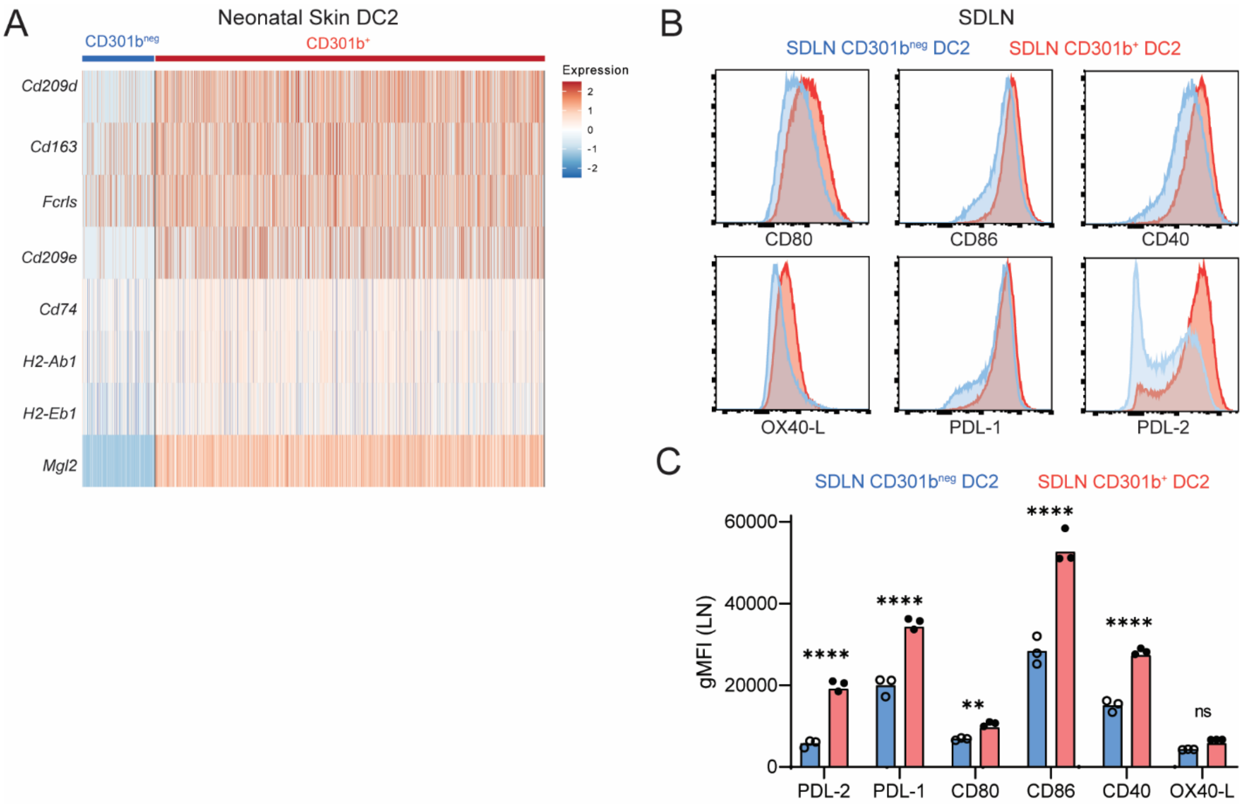


**Figure S4: Related to main Figure 4** (A) Heatmap of the normalized expression of major genes of interest overexpressed in CD301b+ DC2 compared to CD301b$^{neg}$ DC2. (B) and (C) Spectral flow cytometry of activation markers in DC2 from SDLN of D10 mice 16h after *S. epidermidis* skin colonization (B) Representative histograms and (C) quantification of the geometric mean of fluorescent intensity (gMFI) of the indicated markers. Each dot is biological replicate. 1 of 2 independent experiments are shown. Two-way ANOVA with Šídák's multiple comparisons test. . **p < 0.01, ****p<0.0001, ns = not significant.

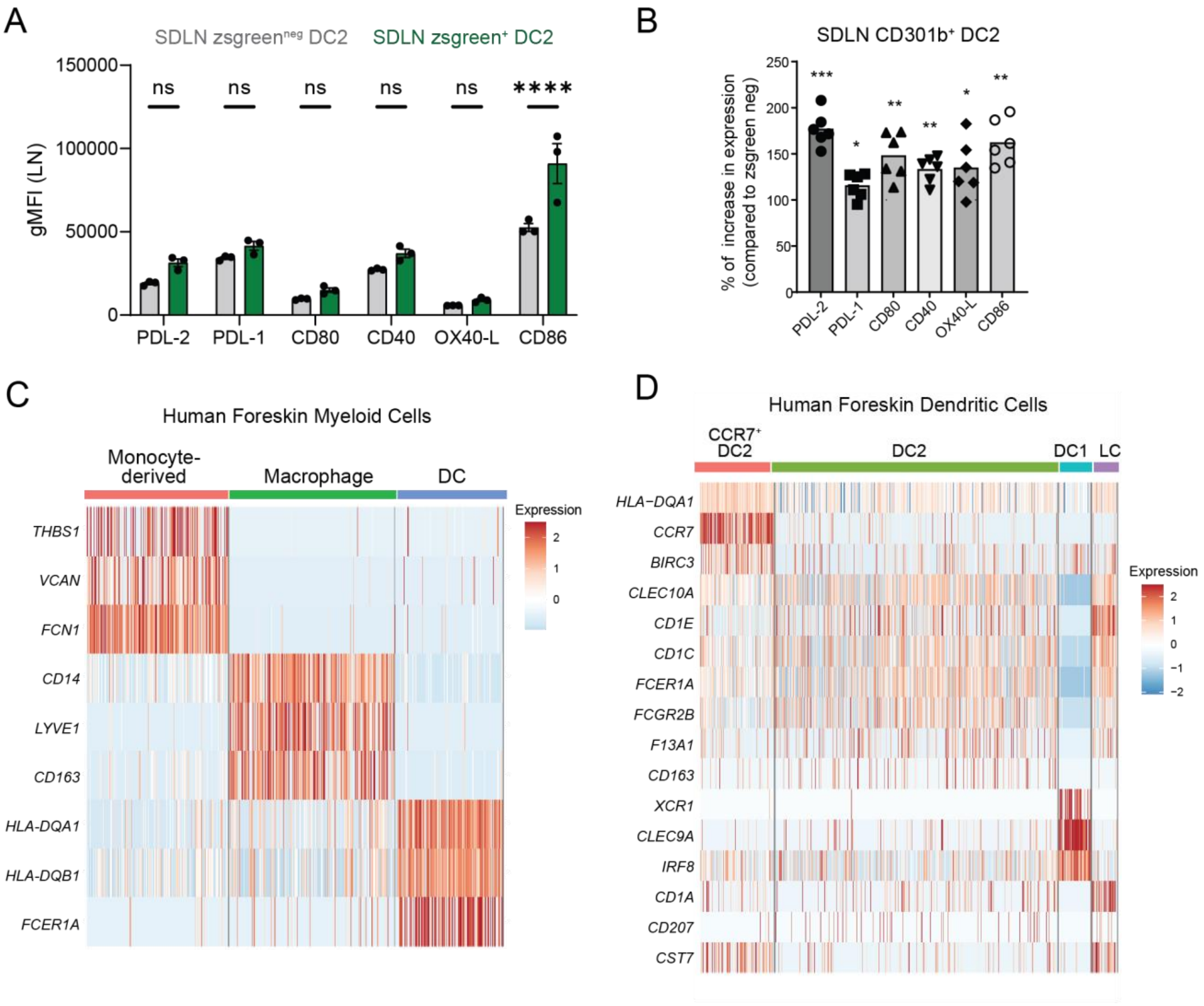


**Figure S5: Related to main Figure 5** (A-B) Spectral flow cytometry comparing protein-level expression of key markers of interest in commensal antigen-loaded (zsgreen$^{+}$) and unloaded (zsgreen$^{neg}$) CD301b$^{+}$ DC2 in neonatal SDLN 16h after *S. epi*-zsgreen colonization. (A) Quantification of the gMFI in loaded vs non loaded CD301b$^{+}$DC2. Each dot is a biological replicate pooled from 3 neonatal mice. 1 out of 2 independent experiments is shown. Two-way ANOVA with Šídák's multiple comparisons test. (B) Representation of the same data as relative expression in zsgreen$^{+}$ vs. zsgreen$^{neg}$ DC2. Data pooled here from two replicate experiments, each data point is a biological replicate pooled from 3 neonatal mice. One sample Wilcoxon test to a hypothetical value of 100. (C) Heatmap of genes used to cluster the human scRNAseq myeloid data in Fig. 5D. (D) Heatmap of genes used to cluster the human neonatal foreskin DC subpopulations as shown in Fig. 5E.

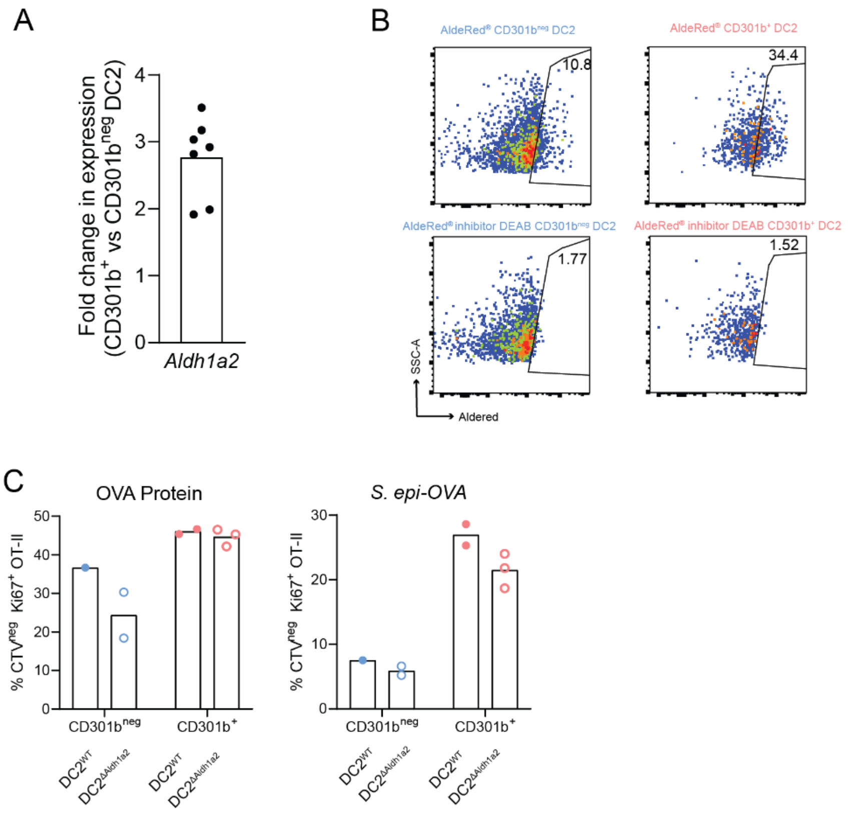


**Figure S6: Related to main Figure 6** (A) *Aldh1a2* expression in CD301b$^{+}$ DC2 vs CD301b$^{neg}$ DC2 from SDLN of D10 pups, as fold-change relative to housekeeping gene. (B) Representative plots of AldeRed fluorescence on the indicated sorted DCs. (C) CD301b$^{+}$ and CD301b$^{neg}$ DC2 from the SDLN of neonatal *Cd11c*$^{cre}$ *Aldh1a2*$^{fl/fl}$ and *Aldh1a2*$^{fl/fl}$ mice were incubated with OVA or *S. epi*-OVA followed by CTV-labelled OT-II in suboptimal Treg inducing conditions. Quantification of OT-II proliferation.
